# Hour-long, Kilohertz Sampling Rate 3D Single-virus Tracking in Live Cells Enabled by StayGold Fluorescent Protein Fusions

**DOI:** 10.1101/2024.03.14.585070

**Authors:** Yuxin Lin, Jack Exell, Haoting Lin, Chen Zhang, Kevin D. Welsher

## Abstract

The viral infection process covers a large range of spatiotemporal scales. Tracking the viral infection process with fluorescent labels over long durations while maintaining a fast sampling rate requires bright and highly photostable labels. StayGold is a recently identified green fluorescent protein that has a greater photostability and higher signal intensity under identical illumination conditions as compared to existing fluorescence protein variants. Here, StayGold protein fusions were used to generate virus-like particles (StayGold-VLPs) to achieve hour-long 3D single-virus tracking (SVT) with one thousand localizations per second (kHz sampling rate) in live cells. The expanded photon budget from StayGold protein fusions prolonged the tracking duration, facilitating a comprehensive study of viral trafficking dynamics with high temporal resolution over long timescales. The development of StayGold-VLPs presents a simple and general VLP labeling strategy for better performance in SVT, enabling exponentially more information to be collected from single trajectories and allowing for the future possibility of observing the whole life cycle of a single virus.

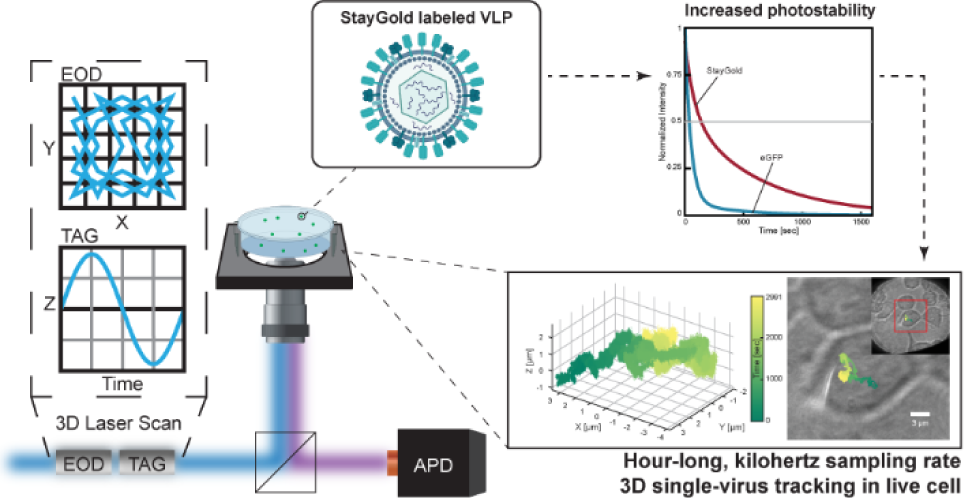

## INTRODUCTION

Over the past few decades, single-particle tracking (SPT) methods have demonstrated their strength in understanding the dynamics of microscopic particles and their interactions within complex environments.^1–7^ SPT methods provide detailed information on the dynamics of single particles and critical molecular subpopulations with high spatiotemporal resolution; information that is obscured by the ensemble-averaged results provided by traditional bulk experiments. As an essential subcategory of SPT, single-virus tracking (SVT) methods have revealed the single viral particle dynamics underlying the complicated mechanisms of viral infection.^8, 9^ Researchers have utilized SVT to capture extracellular viral diffusion,^10, 11^ visualize virus-receptor interactions,^12^ probe viral envelope fusion events,^13^ identify intercellular trafficking mechanisms along the cytoskeleton,^14^ and identify how cell surface structures such as filopodia facilitate viral infection.^15^ While various imaging modalities can be employed for SVT, fluorescence microscopy is widely utilized due to its high sensitivity, versatility, viability, and compatibility with live cell imaging.^16,17^

The crucial steps in viral infection cover a vast range of spatial and temporal scales. A complete viral infection circle lasts from hours to days at the cell and tissue level,^18^ while critical molecular scale steps, such as viral attachment and receptor binding, may happen on the scale of nanometers and milliseconds or even faster.^19^ Given the limited total amount of photons that can be emitted by fluorophores, typically organic dyes or fluorescent proteins (FPs),^10–14, 20, 21^ it is very challenging to observe a single viral particle with high spatiotemporal resolution over a sufficient time period to capture the entire infection process. Fortunately, recent advancements by Hirano and coworkers in 2022 led to the development of the StayGold green fluorescent protein, which is brighter and dramatically more photostable than existing fluorescent proteins.^22^ StayGold forms an obligate dimer with a molecular mass of ∼ 59.9 kDa,^22^ approximately twice as large as eGFP (∼ 26.9 kDa).^23^ It was also reported that StayGold had a comparable maturation rate to the rapidly-maturing mNeonGreen.^22^ In the initial report by Hirano et al., this novel FP showed an increased quantum yield (0.93) compared to eGFP (0.72), ^22^ and comparable or increased quantum yield than the manufacturer-reported quantum yields of organic dyes such as ATTO 488 (0.80),^24^ and Alexa Fluor 488 (0.92).^25^ The increased brightness means more photons are available per unit time, while the increased photostability means more total photons are available for monitoring the protein before irreversible photobleaching. In conventional microscopy, the precision is given by 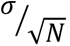, where σ is the size of point spread function and *N* is the number of collected photons. Therefore, the expanded photon budget from StayGold fluorescent proteins has the potential to increase the sampling rate, localization precision, and duration of observation. When applied to SVT, these newly engineered probes provide the possibility of both the high sampling rate and long-term observation required for molecular-scale observation of single viral particles throughout the entire infection process.

In this paper, we engineered Vesicular stomatitis virus G protein (VSV-G) pseudotyped lentiviral particles, packaged with StayGold fluorescent protein viral protein (Vpr) fusions (Vpr-StayGold). The produced virus-like particles (VLPs) were examined and characterized with 3D Single Molecule Active-feedback Real-time Tracking (3D-SMART) microscopy and exhibited extreme resistance to photobleaching as compared to similar constructs. Then, StayGold-VLPs were introduced into the live cell context as a probe to study intracellular viral trafficking. It was demonstrated that VLPs internally labeled with StayGold retained infectivity and significantly prolonged the observation duration of SVT to over an hour, dramatically outlasting eGFP labeled VLPs.^10^ The extended observation period provides extra information and is an important step towards observation of the entire infection process from the perspective of a single virus with high spatiotemporal resolution.

## MATERIALS AND METHODS

### 3D-SMART Overview

High-speed single-viral-particle tracking was accomplished by 3D Single-Molecule Active-feedback Real-time Tracking (3D-SMART) microscopy initially developed by Welsher and co-workers.^26–28^ In this method, a continuous-wave solid-state 488-nm laser source (FCD488-30, JDSU) is used for excitation. The laser beam is deflected by a pair of electro-optical deflectors (EODs. M310A, ConOptics) in a Knight’s tour pattern in the *XY* plane. A tunable acoustic gradient (TAG. TAG 2.0, TAG Optics) lens shifts the laser focus axially and together with the EODs forms the 3D scan volume (1 μm × 1 μm × 2 μm, *XYZ*). The collected photon arrivals are detected with a single-photon counting avalanche photodiode (APD, SPCM-ARQH-25, Excelitas). In real time, the photon arrivals and corresponding laser scan positions are analyzed using an assumed Gaussian density Kalman filter to estimate particle position within the laser scan volume.^29^ An integral feedback controller is used to drive a piezoelectric nanopositioner (Nano-PDQ275HS, Mad City Labs) and a piezoelectric lens positioner (Nano-OP65HS, Mad City Labs) to recenter the particle in the center of the scan volume. The EODs, TAG lens, piezoelectric stage, and APD are fed into a field-programmable gate array (FPGA, NI-7852r, National Instruments).

To contextualize the 3D trajectories synchronously, a brightfield imaging microscope was integrated into the existing 3D-SMART setup. A 780-nm fiber-coupled LED (Thorlabs, no. M780F2) is used as the light source and a high framerate sCMOS camera (pco.edge 4.2, PCO) is used for imaging. The exposure time used is 10 ms and the delay time between each frame is 990 ms for a frame rate of 1 Hz.

The instrumental design is shown in Supporting Information fig. S1.

### Plasmid Construction

The expression vector Vpr-eGFP, used to generate internally eGFP labeled VSV-G VLPs, was constructed as previously described.^10^

The expression vector Vpr-StayGold, used to generate internally StayGold labeled VSV-G VLPs, was constructed as follows. The vector pVpr+ backbone was derived from Vpr- mGreenLantern (see the Supporting Information) and amplified with PCR with the following primers: 5′-AGCGGCCGCGACT-3′ and 5′-GGTGGCGACCGGTG-3′. DNA encoding for StayGold was amplified by PCR with the following primers: 5′- CCGATCCACCGGTCGCCACCGGGGGATCCATGGCC-3′ and 5′- GATCTAGAGTCGCGGCCGCTTTACAGGTGGGCCTCCAGG-3′, using pCSII-EF/mt-(n1)StayGold (Addgene, no. 185823) as the template. PCR products were overlapped and assembled to create Vpr-StayGold with In-Fusion Snap Assembly cloning kits (Takara Bio, no. 638947); successful insertion was verified by sequencing.

Another similar expression vector of Vpr-StayGold containing a shorter 3 amino acids (aa) linker (compared to the 9 aa linker one mentioned above) between Vpr and StayGold was also prepared. However, the plasmid with a shorter linker failed to produce detectable VLPs. These results suggest that an adequately long linker between the StayGold and Vpr DNA may be critical.

### Production of VSV-G VLPs containing Vpr-StayGold or Vpr-eGFP

The pseudotyped lentiviruses with VSV-G spike protein and fluorescently tagged Vpr were produced by transfecting the target cells with corresponding lentiviral plasmids. Before the production of VLPs, 293T/17 cells were grown in completed RPMI-1640 (Millipore Sigma, no. R8758) supplemented with 10% tetracycline-free FBS (Neuromics, no. FBS002-T) and 1 × penicillin-streptomycin (Corning, no. 30-002-CI). The completed RPMI-1640 was also used as the transfection medium and the medium used to collect the virus.

VSV-G VLPs containing Vpr-eGFP and Vpr-StayGold were produced using Lenti-X™ Packaging Single Shots (VSV-G, Takara Bio, no. 631275), with pLVXS-ZsGreen1-Puro (Takara Bio, no. 632677) as the lentiviral vector plasmid DNA and equal amounts of additional Vpr-eGFP or Vpr-StayGold plasmid.

In all VLP preparations above, the cell line used was 293T/17 (Duke Cell Culture Facility, ATCC# CRL-11268). The VLPs produced were all concentrated with Lenti-X™ Concentrator (Takara Bio, no. 631231) and were resuspended with PBS (Genesee Scientific, no. 25-507).

Lentivirus titration was determined by p24 ELISA using Lenti-X™ p24 Rapid Titer Kit (Takara Bio, no. 632200) and a plate reader (PerkinElmer Victor3 V). The lentiviral titers were reported in infectious units per mL (IFU mL^-1^), typically 5-8 × 10^9^ IFU mL^-1^. The VLP stock solutions were stored in aliquots at -80 °C.

### Cell Culture

293T/17 cells (Duke Cell Culture Facility, ATCC# CRL-11268) were grown using complete DMEM, containing DMEM (Corning, no. 10-013-CV) supplemented with 10% FBS (Millipore Sigma, no. F2442) and 1 × penicillin-streptomycin (Corning, no. 30-002-CI) in T12.5 flasks (Genesee Scientific, no. 25-205). Cells were maintained at 37 °C with 5% CO_2_ and passaged when they reached ∼ 70% confluency.

### Microscope Sample Preparation

At 24 h prior to live-cell single-virus tracking experiments, 293T/17 cells were plated in complete DMEM at 8 × 10^5^ cells per well in a 6-well plate with an autoclaved glass coverslip (VWR, no. CLS-1760-025) to ensure 80% confluency on the day of the experiment. The coverslips were coated with poly-L-lysine (Millipore Sigma, no. P6282) for better adhesion of 293T/17 cells and to avoid stacking of cells in multiple layers. Before tracking experiments, the coverslip was transferred to a custom-built sample holder to which HEPES pH 7.4 buffered solution (Live Cell Imaging Solution, LCIS, ThermoFisher, no. A14291DJ) with 2% FBS was added. The desired type of VLPs were added to cells with a final concentration of 1.9 × 10^7^ IFU mL^-1^ (MOI ∼ 3). The VLP treated cells were incubated at 0°C for 30 minutes to allow VLPs to bind to the cell surface while minimizing internalization during incubation. The sample was then positioned onto a heated stage (37 °C. Heating elements: Bioscience Tools, TC-E35. Temperature controller: Bioscience Tools, TC-1-100). All experiments were performed using live 293T/17 cells without staining.

The average excitation power at the objective focus was 22 nW and 7.5 μW for the 488-nm laser beam and 780-nm LED beams, respectively. The exposure time of the sCMOS camera was 10 ms and delay time between each frame is 990 ms to make the imaging frame rate 1 Hz. All the tracking experiments were performed at room temperature (∼ 23 °C).

### Sliding Window Drift Velocity Calculation

The trafficking of individual VLPs in live cells was characterized by anomalous diffusion, with diffusion on short timescales and a net displacement or “drift” on longer timescales. On short timescales, this “drift velocity” can be obscured by the motion due to Brownian diffusion. To quantify the drift velocity, and distinguish it from random thermal motions, a sliding window drift velocity calculation was used.^30^

Assuming the particles undergo Brownian motion in each direction along *X*, *Y* and *Z* axis, *X_i_* = *X*(*iτ*) is denoted as the location of the particle along one-dimensional Brownian path at time *iτ* with *i* = 1, 2, 3 … *M*. The increments between locations are given by *x_i_* = *X_i_*_+1_ - *X_i_* . For Brownian motion, the *x_i_* are normally distributed with mean *uτ* and variance 2*Dτ*, where *u* is the drift velocity and *D* is the diffusion coefficient. That is to say, the drift velocity in each dimension is calculated as:

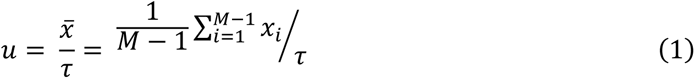

In the analysis, the time interval between each point (*τ*) is 1 second, the window time (*M*) is 200 seconds, and the window step size is 1 second. The large time interval and window time is selected due to the relatively slow motion of viral particles undergoing intercellular trafficking. The drift velocity along the *X*, *Y*, and *Z* axis (*u_x_*,*u*_y_, and *u_z_*) are calculated independently. The magnitude of the drift velocity in the *XY* plane is calculated as

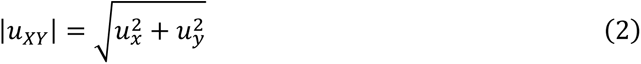

and the direction *θ* of the drift in the *XY* plane is given by

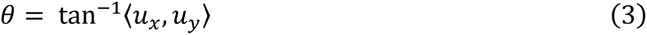

### Spherical Surface Fitting

Particle diffusion on a sphere was fit to a spherical surface using a custom simplex search algorithm. The segment was initially sampled at 10 Hz to obtain a discrete point cloud, though this sampling rate can be varied depending on the size of the data set. The loss function was defined as the residual of the sum of squares between the distance of a spot relative to surface of a given sphere with center of (*x*, *y*, *z*) and radius *r* . For a given center (*x*, *y*, *z*) of a sphere, the optimal *r* is the average distance from all points to the center. As a result, only 3 parameters are needed to find the center of the sphere in the cartesian system. A simplex search MATLAB script using the Nelder-Mead algorithm developed by Singer and Nelder was implemented with an initial simplex size of 3 × 3 × 3 to a 3D grid with each spot 50 nm away from its nearest neighbor was used to find the center of center of sphere.^31^

### Great-Circle Distance Calculation in Spherical Coordinates System

We denote *θ*_1_, *φ*_1_ and *θ*_2_, *φ*_2_ as the geographical longitude and latitude of two points 1 and 2, Δ*θ,* Δ*φ* as their absolute differences, and Δ*σ* as the central angle between them. Due to the relatively small distances compared to the spherical radius, the haversine formula was used to calculate Δ*σ* , the central angle between point 1 and point 2 as:

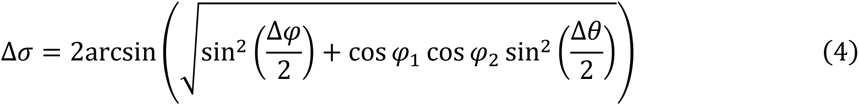

Given the angles in radians, the actual arc length *d* on a sphere of radius *r* can be trivially computed as:

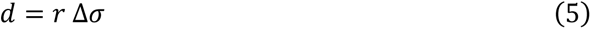

## RESULTS AND DISCUSSION

### Preparation of VLPs labeled with eGFP and StayGold fluorescent proteins

Vesicular stomatitis virus (VSV)-G-pseudotyped VLPs were chosen as the model for evaluation of StayGold labeling due to its ubiquitous host entry receptor, the low-density lipoprotein receptor (LDLR),^32^ resulting broad cellular tropism to infect multiple cell lines. An efficient and well-characterized internal virus labeling approach was utilized whereby fluorescent proteins were fused to the HIV-1 Vpr and packaged into budding lentiviruses.^33^ Two types of different transfer plasmids were used to produce VLPs packaging Vpr-StayGold and Vpr-eGFP (StayGold-VLPs and eGFP-VLPs, Fig. 1a). An immunofluorescence assay was performed to confirm the successful incorporation of Vpr-StayGold or Vpr-eGFP into single virions with the desired spike protein (Fig. 1b-c, fig. S2). An infection assay indicated that the presence of Vpr-StayGold or Vpr-eGFP does not prevent gene delivery (fig. S3).^34^ During the preparation of Vpr-StayGold plasmids, it was observed that short linkers led to poor expression level, as no fluorescent VLPs were observed after production (data not shown). A plausible explanation was that the dimeric nature of StayGold required more space for proper folding thus longer linkers were favored.

**Figure 1.**
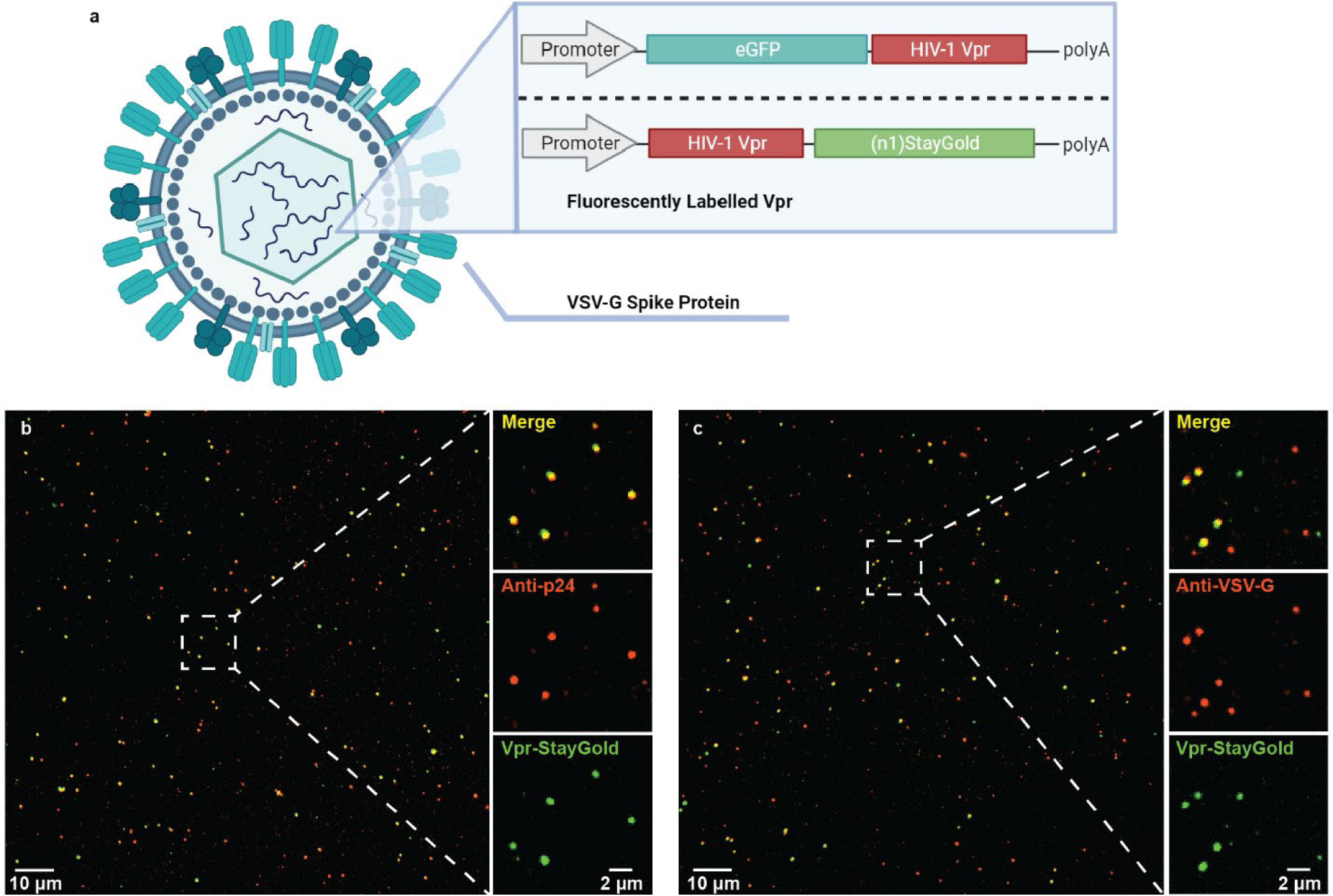
(**a**) Schematic diagram of VSV-G lentiviral particles containing fluoroscently labelled Vpr. The insert shows domain structures of two types of transfer plasmids used to produce the lentiviruses, Vpr-eGFP and Vpr-StayGold respectively. (**b**) A representative image of anti-p24 immunofluoroscence of VSV-G VLPs packaging Vpr-StayGold incubated with primary and secondary antibodies. False-color merge of Vpr-StayGold (green) and AF555 labelled p24 (red). A zoom-in view is shown in right panel. Top: Merge of two channels below. Middle: Channel of AF555 labelled p24. Bottom: Channel of Vpr-StayGold. (**c**) A representative image of anti-VSV-G immunofluoroscence of VSV-G VLPs packaging Vpr-StayGold incubated with primary and secondary antibodies. False-color merge of Vpr-StayGold (green) and AF555 labelled VSV-G (red). A zoom-in view is shown in right panel. Top: Merge of two channels below. Middle: Channel of AF555 labelled VSV-G. Bottom: Channel of Vpr-StayGold.

### High spatiotemporal SVT via 3D-SMART

3D-SMART microscopy was employed to capture viral dynamics with high spatiotemporal resolution (Figure 2). A pair of EODs together with a TAG lens were utilized to generate a rapid 3D scanning volume with the size of 1 μm × 1 μm × 2 μm in *XYZ*, with scan frequencies of 50 and 70 kHz in *XY* and *Z*, respectively. The collected photon arrivals were captured by an APD and were used every 20 μs to estimate the real-time position of the particle within the scan volume. Feeding these estimated positions into an integral feedback control loop, a piezoelectric stage was then employed to recenter the particle within the 3D scan volume as it moves. The tracked particle is maintained within the center of the laser scan volume throughout the duration of the trajectory. As a result, the readout of the piezoelectric nanopositioners can be used as a measure of particle position. By this method, 3D-SMART can produce 3D particle trajectories with 1 kilohertz sampling rate, limited by the 1-ms mechanical response time of the stage, and with localization precision down to ∼ 20 nm in *XY* and ∼ 80 nm in *Z* (previously reported in Ref. ^10^). This sampling rate can be improved by improving the response time of the feedback element. By replacing the *XY* piezoelectric stage with a galvo scanning mirror, the sampling rate could reach approximate 5 kHz.^35^ This sampling rate can theoretically be improved further. In 3D-SMART, since a single-photon counting APD is used, the position estimate of the particle can be updated with each arriving photon. For the particles described below, which have emission intensities in the 50 to 100 kHz range, the position could be updated every 10 to 20 µs, though with an intrinsic trade-off between temporal resolution and localization precision. If there were infinite amount of photons, the bin time could be further narrowed down and the temporal resolution would only be ultimately limited to the photon collection rate.^36^ The additional photons provided by the StayGold fluorescent proteins are critical to increasing both the observation window and sampling rate in SVT.

**Figure 2.**
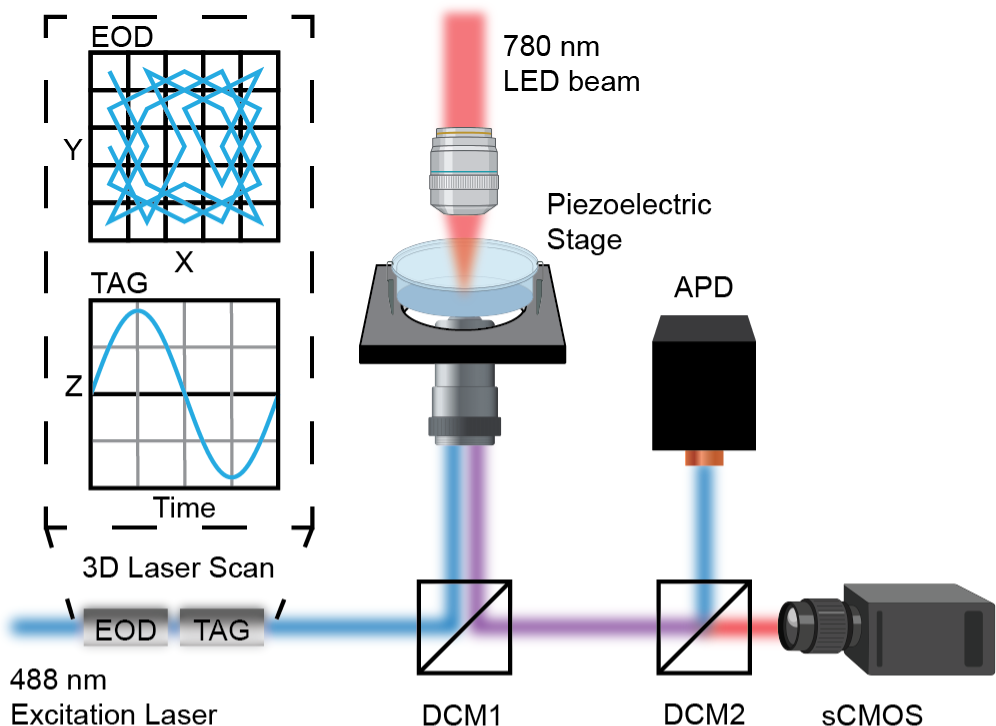
Illustration of 3D-SMART integrated with brightfield imaging. A 488-nm laser is used for tracking excitation. It is deflected by a pair of electro-optical deflectors (EODs) and a tunable acoustic gradient (TAG) lens to form a 3D scanning volume. The beam is reflected into the objective by a multiband dichroic mirror (DCM1). A 780 nm LED beam is used for the brightfield imaging excitation and shines directly from the top of the sample. The combined imaging and tracking emission signal were then sepreated by a dichroic mirror (DCM2). The imaging signal was detected by a sCMOS camera and the tracking signal was detected by a single-photon counting avalanche photodiode (APD).

To provide label-free imaging of the surrounding cellular environment, a brightfield imaging setup was integrated with the 3D-SMART microscope for hour-long contextualization of 3D viral trajectories with kHz sampling rate (Fig. 2). The label free imaging helps maintain the viability of the cells and helps reduce labeling induced artifacts that may be misconstrued as virus-cell interactions.^37^ The imaging setup consists of a 780 nm LED and a high framerate sCMOS camera. Calibration was performed to register 3D-SMART trajectories with the brightfield images to generate the final results (fig. S4-5).

### Characterization of VLPs labeled with eGFP and StayGold fluorescent proteins

Using 3D-SMART, kHz-rate 3D-SVT was performed on VLPs freely diffusing in LCIS at room temperature for solution-phase single-VLP characterization. Particle size was calculated by mean square displacement (MSD) analysis and hydrodynamic size was extracted via the Stokes-Einstein equation. Particle brightness was evaluated by the mean intensity of the first second of each trajectory (Fig. 3a). A total number of 98 for StayGold-VLPs and 52 for eGFP-VLPs trajectories were used to yield the following characterization. The produced StayGold-VLPs (180 ± 72 nm) were approximately 50% larger than eGFP-VLPs (120 ± 47 nm, Fig. 3c). Following a similar trend, StayGold-VLPs (90.51 ± 38.78 kHz) were roughly twice as bright as eGFP-VLPs (48.50 ± 19.45 kHz, Fig. 3d). The differences in size and intensity of two VLPs could be caused by different amount of Vpr packed inside, which could be estimated by the emission intensity of single Vpr- FPs. Measurements were carried out to determine the emission intensity of a single Vpr-eGFP (∼ 644 Hz) or Vpr-StayGold (∼ 872 Hz) via single molecule photobleaching (fig. S6). Based on the intensity of single FPs and the ensembled intensity of VLPs, it can be determined that 40% more Vpr-StayGold (104 ± 44) were packaged in a single VLP as compared to Vpr-eGFP (74 ± 30), which likely contributed to the size and intensity difference. It is expected that the increase in size of the StayGold VLPs relative to the eGFP-VLPs is driven by increased packaging of Vpr within the capsid. This could be adjusted in future works to reduce this size difference. For the current work, the sizes of both eGFP-VLPs and StayGold-VLPs fell in the typical range of 100-200 nm scene in other reports. ^38, 39^

**Figure 3.**
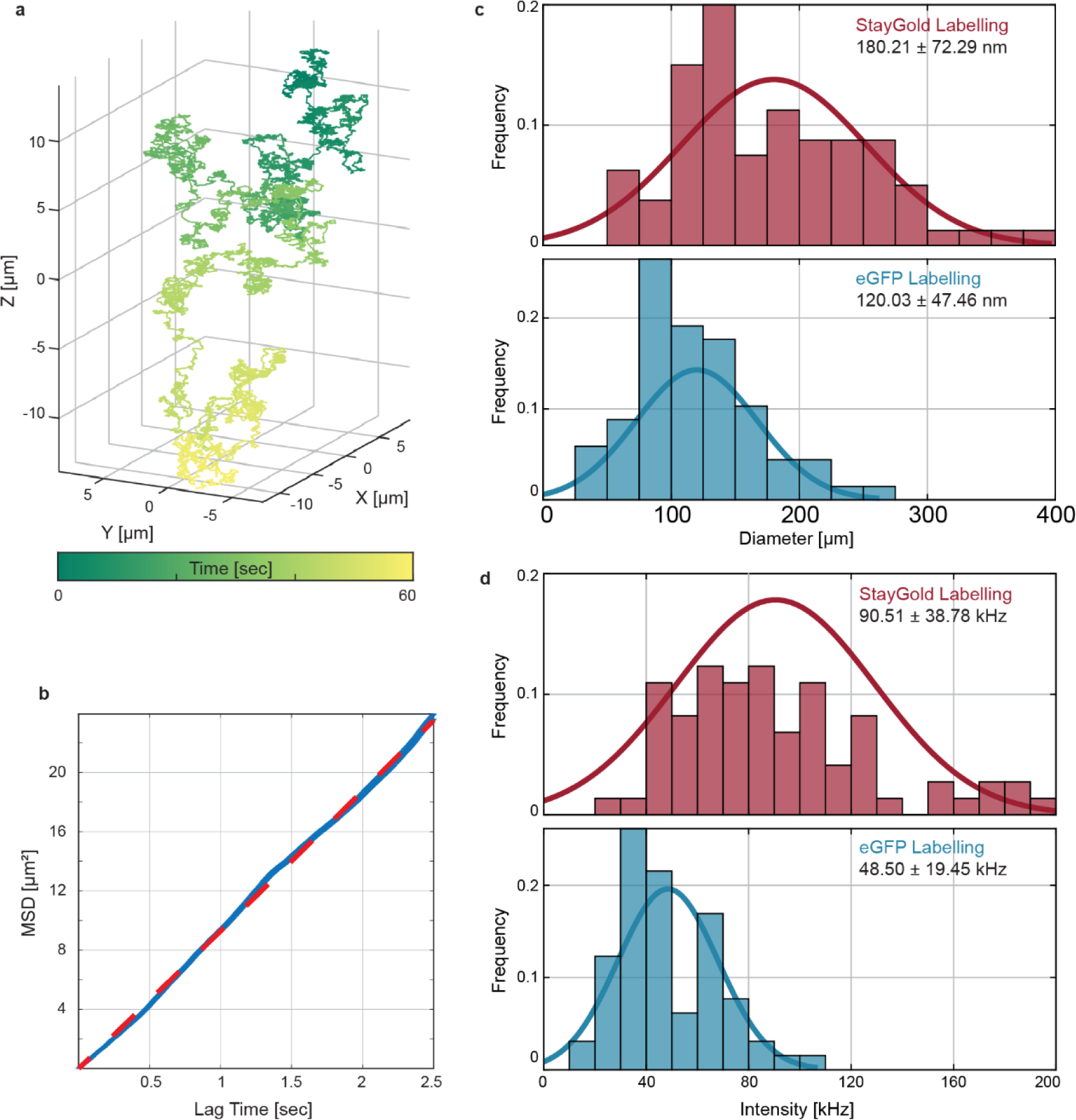
Solution-phase single-particle characterization of StayGold-VLPs (N = 98) and eGFP-VLPs (N = 52) . (**a**) Example 3D trajectory of a single StayGold-VLP obtained by 3D-SMART. (**b**) MSD analysis of (**a**). The slope of linear fit (red) from MSD plot (blue) yielded diffusion coefficient and particle size could be calculated based on Stoke-Einstein equation. (**c**) Size distribution of StayGold-VLPs (upper, red) and eGFP-VLPs (lower, blue). (**d**) Intensity distribution of StayGold-VLPs (upper, red) and eGFP-VLPs (lower, blue). For the free VLP tracking experiments, the average excitation power at the objective focus was ∼ 43 nW. The curves in (**c**) and (**d**) are normal distribution with the same mean and standard deviation of VLPs’ size and intensity, respectively.

The starkest difference between the two types of VLPs was revealed when their photostability was investigated. To measure the photostability, surface-immobilized StayGold-VLPs and eGFP- VLPs were illuminated until photobleached to the background level (Fig. 4a). The photobleaching curves of StayGold-VLPs were well fitted to bi-exponential decay model, which indicated that bleaching may occur through multiple pathways,^40^ and the corresponding half-lives under different excitation powers were calculated according to the fitting. Under the same ∼ 85 nW excitation power at the sample, StayGold-VLPs displayed about four-fold longer half-life time than eGFP- VLPs (StayGold-VLPs: 166 ± 18 sec vs. eGFP-VLPs: 40 ± 7 sec, 95% prediction bound, Fig. 4b). In the range of our tested excitation power, the lifetimes of StayGold-VLPs and eGFP-VLPs also show a similar power dependence to that reported in Ref. 22 (Fig. 4c). The remarkable photostability of the StayGold-VLPs indicated that it is a promising labeling method for SVT that could enable information on viral dynamics on timescales currently inaccessible by existing labeling methods.

**Figure 4.**
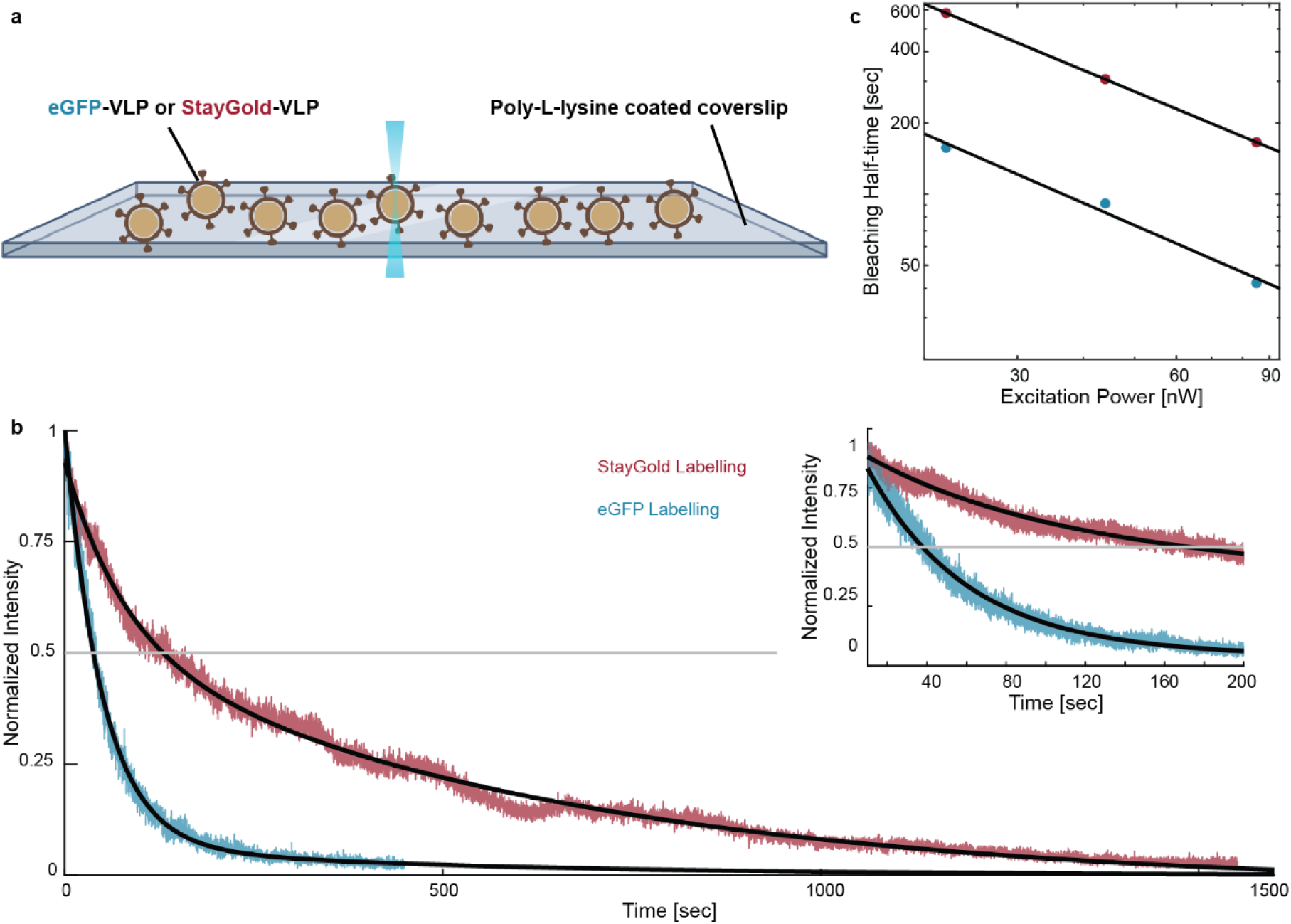
Photobleaching experiments of StayGold-VLPs and eGFP-VLPs. (**a**) Illustration of photobleaching experiments. (**b**) Photobleaching curve of StayGold-VLPs (red) and eGFP-VLPs (blue) showed significantly improved photostablity of StayGold labelling. The overlayed black lines corresponded to the fitting results. Photobleaching data for StayGold-VLPs were fitted to the bi-exponential decay equation 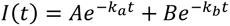. *A* = 0.39, *k_a_* = 0.013, *B* = 0.54, *k_b_* = 0.0016. Photobleaching data for eGFP-VLPs were fitted to the mono-exponential decay equation *I*(*t*) = *Ae*^−*kt*^. *A* = 0.95, *k* = 0.016. The average excitation power at the objective focus was ∼ 85 nW in the photobleaching experiment. Insert: zoom in of the photobleaching curve in (**b**). (**c**) log-log plot of bleaching half-time of StayGold-VLPs (red) and eGFP-VLPs (blue) and excitation power showed the power dependance. Data were fitted to the equation log(*Y*) = −αlog(*X*) + *c*. *α* value was 0.93 and 0.97 for StayGold-VLPs and eGFP-VLPs, respectively, which were close to previously reported value 0.90 and 0.97 in Ref. 22.

### Hour-long, kilohertz sampling rate SVT of VLP-StayGold in cells

The performance of StayGold-VLPs for SVT was validated with live-cell experiments. StayGold-VLPs were incubated with live cells at 4℃ for 30 minutes to allow VLPs to bind to the cell surface. The VLP treated cells were then transferred to the 3D-SMART microscope and heated to 37℃, which was maintained throughout data collection. The live-cell context of 293T/17 cells was recorded with the brightfield imaging approach detailed above. To further prolong the tracking duration, the excitation power was reduced to ∼ 22 nW at the laser focus.

Control experiments were performed to ensure that the particles being tracked within live 293T/17 cells were in fact VLPs, and not just autofluorescence puncta (fig. S7). Trajectories of autofluorescent puncta, which were observed in cells without exposure to fluorescent VLPs, had much lower pickup intensity (initial intensity when tracking was engaged) and bleached rapidly compared to the longer lasting fluorescent VLPs. The pickup intensity of ∼ 97% of autofluorescent trajectories were below 20 kHz (16.76 ± 2.17 kHz) while over 76% of trajectories with fluorescent VLPs displayed higher pickup intensities (fig. S7a). The tracking duration was less than 100 seconds for 86% of tracked autofluorescent puncta, which was significantly shorter than trajectories of fluorescent VLPs (fig. S7b-c). Based on the pickup intensities and tracking durations, 18 out of 77 trajectories were differentiated as autofluorescent trajectories and were filtered out with 95% confidence, forming a dataset of 59 live-cell StayGold-VLP trajectories (fig. S7d).

For the sake of experimental efficiency, tracking was automatically disengaged when a preset maximum tracking duration was reached, which was set to 4000 seconds (4×10^6^ localizations). Among 59 datasets obtained from tracking StayGold-VLPs, the median tracking duration is ∼ 28 minutes, while most trajectories of eGFP-VLPs lasted less than 10 minutes. Noticeably, 46% of StayGold-VLP trajectories were over half an hour, and 15% of trajectories were over an hour long and 7 trajectories hit the set maximum (fig. S8). It is worth noting there that photobleaching is not the only limitation to trajectory duration. Diffusing particles could exceed the working range of the piezoelectric stage over the long tracking duration, leading to trajectory termination. Trajectories could also end if another nearby VLP was detected by the tracking system. Upon finding a brighter particle, the feedback algorithm will “jump” to this new particle, prematurely ending observation of the initial particle. These “jumps”’ usually happened towards the end of a tracking period due to the decay of particle intensity over time, and pose a further challenge to long duration active-feedback SVT.

Extremely long duration trajectories were routinely observed and demonstrate the immense amount of information that can be obtained by combining StayGold VLPs with 3D-SMART (Fig. 5 and Supplementary Video 1; See Supporting Information for more examples). The example trajectory shown in Fig. 5a lasts for over 3000 seconds (3×10^6^ 3D localizations), in contrast to a typical eGFP-VLP trajectory with an even higher starting intensity (Fig. 5c) which could only be tracked for ∼ 14 minutes before being totally photobleached (reaching the background intensity level). The combination of high sampling rate and long duration observation allows a more complete observation of single viral dynamics across multiple timescales. During a significant length of the observation window of the StayGold VLP shown in Fig. 5a, the viral diffusion was relatively confined with an average drift velocity of < 5 nm/sec. However, at ∼ 1100 sec of the trajectory, a burst in viral motion was observed and the drift velocity increased to a maximum of ∼ 20 nm/sec for 300 seconds (Fig. 5d-f). The prolonged tracking duration from StayGold fluorescent protein fusions made it possible for more successful observation of different viral diffusion patterns, especially those viral particles with dormant periods that are longer than the lifetime of other fluorescent labels.

**Figure 5.**
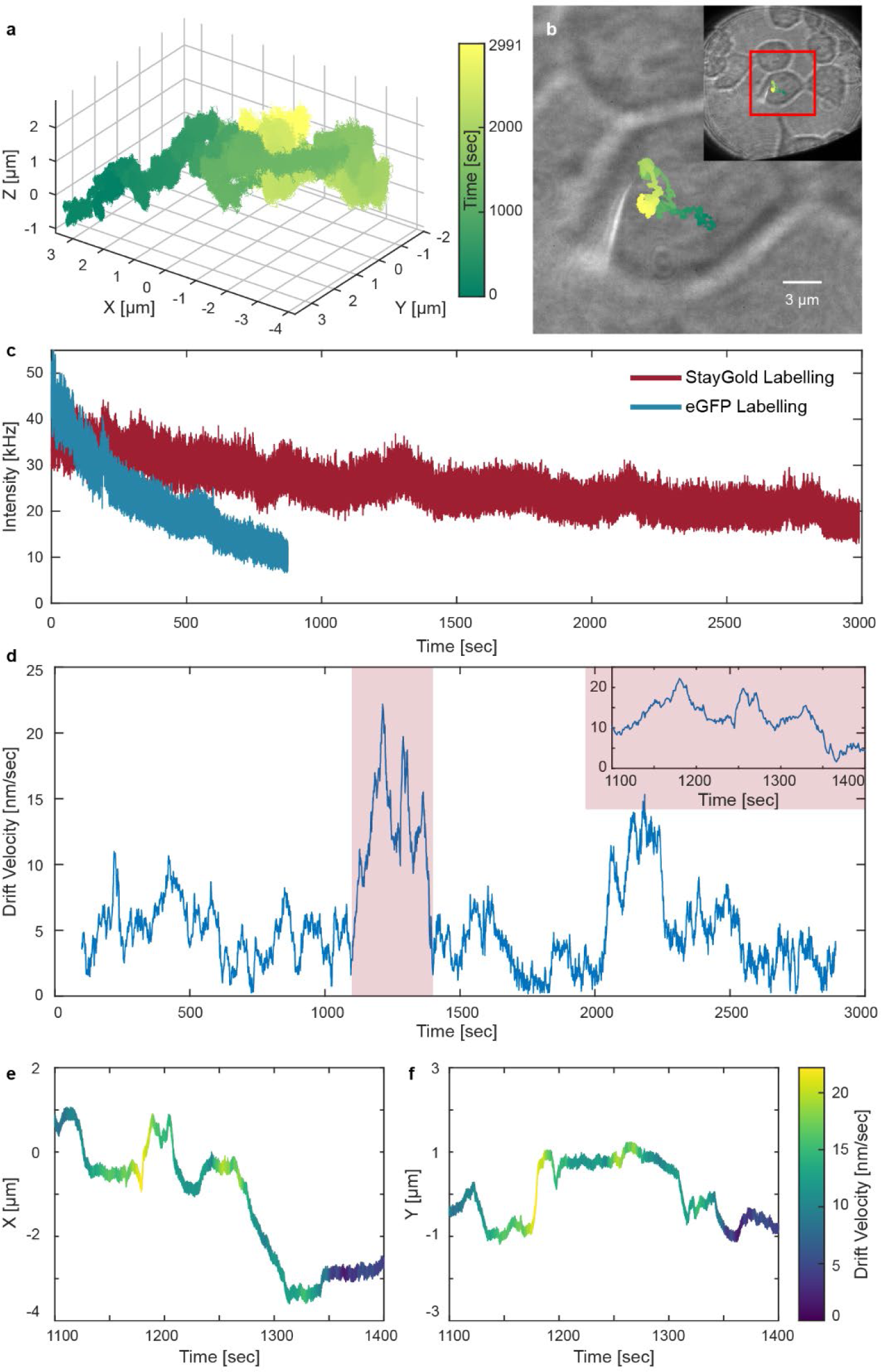
An example of tracking VSV-G Vpr-StayGold lentivirus in live 293T/17 cells (see also Supplementary Video 1, see fig. S9 for another examples). (**a**) 3D trajectory of a StayGold-VLP in a live 293T/17 cell. (**b**) bright field image overlapped of the top-down view trajectory in (**a**). Insert: larger view of cell. The trajectories in (**a**-**b**) are color-coded by time with the same colormap in (**b**). (**c**) Intensity trace of (**a**) and a trajectory of a eGFP-VLP in a 293T/17 cell (see fig. S10 for complete information) showed better photostability and longer observation time benefited from StayGold labelling. (**d**) 2D drift velocity trace of (**a**). (**e-f**) *X* and *Y* trace of (**a**) color-coded by 2D drift velocity with the same colormap in (**f**).

Linear trafficking and membrane diffusion were two major anomalous viral motions observed among the StayGold-VLP trajectories. These two different patterns are easily differentiated using power-law MSD analysis, where the exponent α can distinguish between confined, diffusive, and direction motions (fig. S13). Linear trafficking is characterized by super-diffusion with α > 1 (fig. S13a). In contrast, membrane diffusion was mostly sub-diffusive with α < 1 (fig. S13b). Different values of the power law exponent α did not necessarily correlate with faster or slower diffusion. For example, linear-trafficking segments (fig. S13b) often show a small diffusion coefficient (< 0.1 μm^2^/sec), but a high power-law coefficient (α > 1.5). It was also observed that linear trafficking events were highly anisotropic and directional (fig. S13d).

### Reconstructing 3D surfaces in live cells with long timescale viral trajectories

The long timescale of trajectories enabled by StayGold labeling allowed both kHz-rate 3D localization as well as the carving out of cellular structures within intracellular viral trajectories. The long timescale trajectories traced out the shape of physical cellular structures such as the cytoskeleton and membrane protrusion, demonstrating that SVT is also a tool for studying intracellular structures that play a role in viral infection.^41^ Previous research using 3D-SPT revealed cylindrical patterned trajectories along cell surface microspikes, hemispherical diffusion on membrane blebs, and linear motions along a membrane protrusion.^10, 36, 42^ In the highly-sampled trajectories reported here, directed linear motion could be observed, consistent with viral trafficking along the microtubule network. Uniquely, the extremely long timescale of observation here, combined with high-speed 3D sampling, allowed the large scale geometry of the underlying network to be resolved. An example of such a trajectory is shown in Fig. 6 and fig. S11, where the linear trafficking clearly traced out a curved surface containing the linear motions over a large axial range of about 8 μm. A spherical fit of ∼ 30 minutes of the total trajectory enables extraction of the properties of the underlying structure (Fig. 6a and Supplementary Video 2). Once the 3D structure over which the microtubule network is measured, more accurate calculations of the directed motions of single VLPs along these tracks can be obtained. This is done by converting the original Cartesian coordinates into spherical coordinates on the surface of the larger structure (Fig. 6b and fig. S12a-b).^43^ If a 2D imaging method were used, and directed motion were calculated based on an XY-projection, the drift velocity of viral particles would be dramatically underestimated, as shown in Fig. 6c. A drift velocity analysis of viral motion is far more accurate upon change of coordinates of the local structure. The calculated drift velocity on the fitted spherical surface was significantly larger than just calculating from the *XY*-projection and matched the reported range viral trafficking velocity of about 1-10 μm/min (Fig. 6c).^44, 45^ The reconstruction is possible due to the long timescale observation of the viral motion enabled by StayGold, and would be impossible to glean from shorter duration observation. If the trajectory length is reduced to 400 seconds (fig. S12c), corresponding to the typical lifetime of eGFP-VLPs, key structural information is lost, and the curved motion appears to be close to linear motion on a planar surface. Understanding the nature of the spherical microtubule network traced by the VLPs will require direct imaging of the network and trafficking proteins that participate in the process. The current bright-field imaging approach does not provide this information. Future work will focus on incorporating volumetric imaging with these long-time-scale trajectories to fully understand the underlying biology.

**Figure 6.**
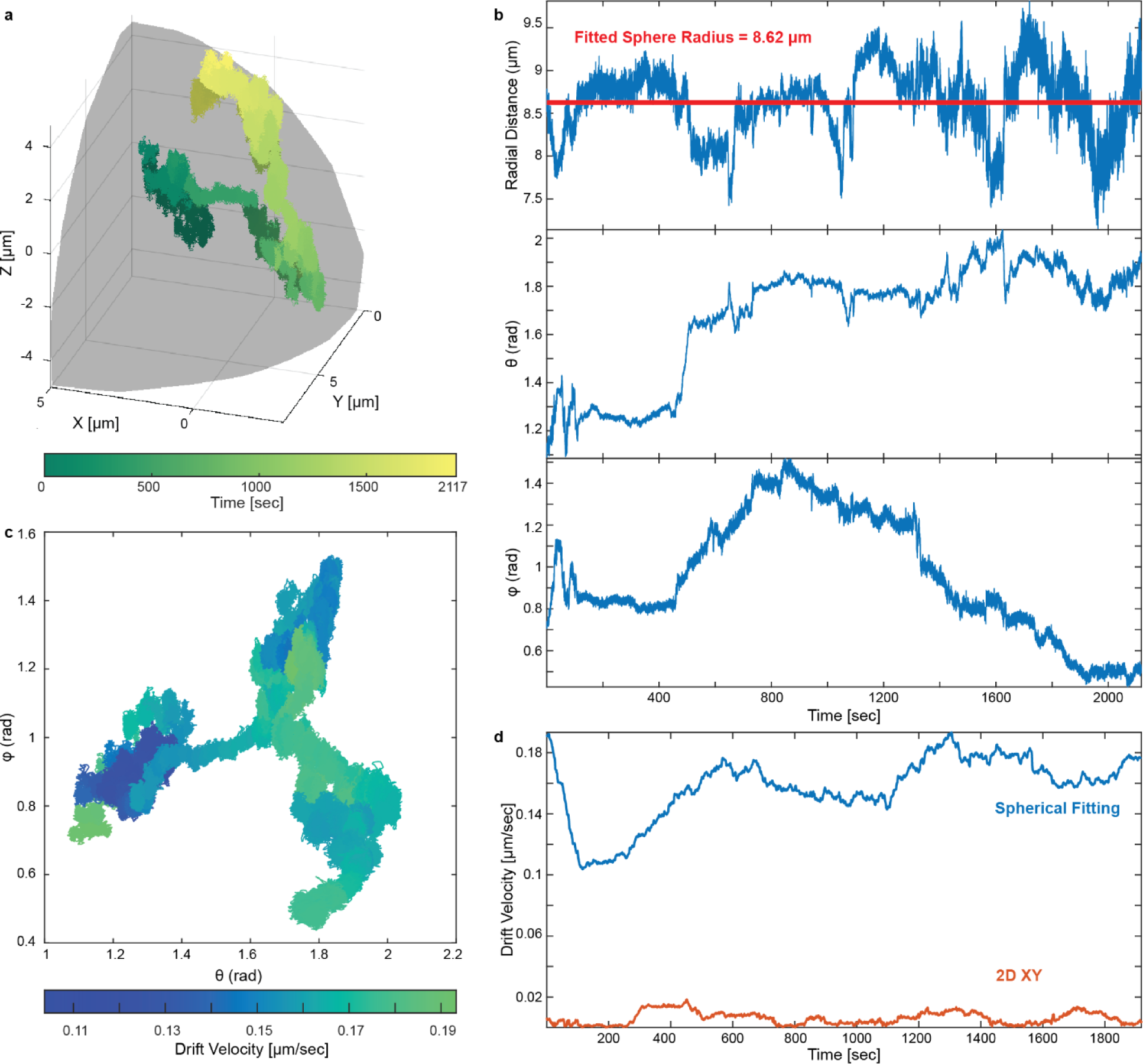
Mapping the viral trafficking trajectory on a spherical surface (see also Supplementary Video 2). (**a**) The segment of trajectory (see fig. S11 for complete information) overlayed with the fitted sphere in cartesian coordinate. The segment was color-coded by time. (**b**) Trace plot of radial distance *r*, polar angle (or longitude) *θ* and azimuthal angle (or latitude) *φ* respectively. The red line on the top plot represents the radius of fitted sphere. (**c**) The spherical surface was projected to a cylinder developable surface then unfolded into a 2D plane. The trajectory was color-coded by drift velocity calculated from spherical fitting. (**d**) Drift velocity trace calculated from the spherical fitting (blue) and *XY* projection (orange) respectively.

## CONCLUSIONS

Despite the assistance of highly photostable StayGold fluorescent proteins, the expanded photon budget remains finite. Thus, it’s prudent to explore methods for utilizing these surplus photons more efficiently. Previous analyses of 3D-SMART have demonstrated that different EOD scanning patterns can affect both localization precision and photobleaching rate.^46^ When a tracked particle moves slowly, it will be tightly locked in the center of the scanning volume. For a well- centered particle, the photons collected from the center pixel in the laser scan provides little to no location information, but still contributes to photobleaching. Instead of using the current Knight’s Tour pattern, switching to a four-corner off-centered pattern could achieve a ∼ 20% increase in precision while reducing the photobleaching rate by avoiding unnecessary scanning of the particle center.^46^ It is also potentially possible to further reduce the excitation laser power to prolong the tracking duration while extending the localization bin time to compensate for decreased signal-to- noise.

Further improvements to the contextualization of these long trajectories are the next step forward. Here, it was difficult to use fluorescent labeling of cellular structures as they bleach on much faster timescales than the StayGold VLPs. The challenge will be to integrate these trajectories with images,^10^ but with the images poled at an infrequent rate to maintain fluorescence from the cells over the hour-long observation time.

Future work will also benefit from the recent development of multiple monomeric StayGold fluorescent proteins by different groups,^47–49^ while maintaining the brightness and photostability of the dimeric variant used here. These monomeric variants should continue to broaden the application scope of these unique labels, while helping to minimize fusion-induced artifacts that can result from multimeric fusions.

In summary, this work demonstrates that VLPs labeled with fluorescent protein fusions of StayGold dramatically extended the tracking duration of kHz 3D-SVT from several minutes to up to an hour. Extending the observation time to the hour timescale with kilohertz sampling rate not only brings exponentially more information but also it is one step closer towards observing the entire infection process of a single virus with molecular-scale precision. The outstanding spatiotemporal resolution in 3D provided by 3D-SMART brought further insights of motions in three-dimensions that are missing or inaccurately quantified in 2D-SVT methods. Moreover, this brighter and more photostable StayGold labeling method is also seamlessly and universally transplantable to other fluorescent methods and applications.

## ASSOCIATED CONTENT

### Supporting Information

The Supporting Information is available free of charge.

Additional information of the instrumental setup, biological sample preparation, and analytical methods. (PDF)

3D single virus tracking trajectory of StayGold-VLP intracellular dynamics (MP4)

3D single virus tracking trajectory of StayGold-VLP diffusion on curved cellular surface (MP4)

## Author Contributions

J.E., Y.L. and K.D.W. conceptualized the study. J.E., Y.L. and K.D.W. were responsible for the methodology. Y.L., J.E. and H.L. performed investigations. K.D.W. was responsible for funding acquisition. Y.L., J.E., H.L., C.Z. and K.D.W. curated the data. Y.L., C.Z. and K.D.W. performed the visualization. Y.L. and K.D.W. wrote the original draft and reviewed and edited the manuscript. The manuscript was written through contributions of all authors. All authors have given approval to the final version of the manuscript.

## Notes

The authors declare no competing financial interest.

## ACKNOWLEDGMENT

We acknowledge financial support from the National Institute of General Medical Sciences of the National Institutes of Health under award number R35GM124868 (K.D.W.). We appreciate the support from the Fitzpatrick Institute for Photonic at Duke University, along with the John T. Chambers Scholarship (to Y.L.). We also acknowledge the Duke Cell Culture Facility for access to cell lines used in this study. We gratefully acknowledge the Duke Light Microscopy Core Facility for their support and assistance in this work. We also appreciate the Franz Lab at Duke University for the access to their plate reader. Some schematics were created with BioRender.com.

