## Supporting Information for "Hour-long, Kilohertz Sampling Rate 3D Single-virus Tracking in Live Cells Enabled by StayGold Fluorescent Protein Fusions"

**This PDF file includes:**

Materials and Methods
Figures S1 to S13
Captions for Movies S1 to S2

**Other Supplementary Materials for this manuscript include the following:**

Movies S1 to S2

### 1 Materials and Methods

#### 1.1 Tracking and Imaging Integration Overview

The 3D-SMART active-feedback tracking and brightfield imaging are coupled into a single platform through the chromatically separated excitation and emission pathways. The stage coordinates of piezoelectric stage define the physical space and can be used as an absolute reference between tracking and imaging spaces. As the 3D-SMART system natively outputs the trajectory in stage-space coordinates, only the location of the brightfield image must be adjusted to generate fully registered tracking and imaging spaces. Multiple calibrations were used to obtain the imaging parameters needed to convert pixel locations from image-space into physical stage-space and are described below.

##### 1.1.1 Instrumental Set-up Overview

Detailed instrumental diagram was depicted in fig. S1. The imaging and tracking emission are merged from the objective and are reflected to the detection pass with a multiband entrance dichroic (**DCM1**). The beam was then split by a 640 nm short-pass dichroic mirror (**DCM2**). A 700 nm long-pass filter (**f1**, Thorlabs, no. FELH0700) is employed to remove undesired signal and the emission filter for the tracking signal detection is a 535/50 bandpass filter (**f2**, Semrock, no. FF01-535/50-25).

##### 1.1.2 Tracking and Imaging Relative Center Positions

The lateral center of the tracking volume sits at a constant location relative to the image, determined by the relative alignment of the tracking excitation beams and camera. The track center is known as described below.

fig. S2 shows the process used to register the relative lateral location between the tracking volume in the image space. The emission filter in front of sCMOS camera was removed and 488-nm tracking laser was turned up to over 1μW to allow enough signal pass through the DCM. 2D Imaging data (fig. S2a) of the excitation beam are acquired for ~20 frames. We assume that the position of the beam corresponds to the lateral center of the tracking volume, and apply a Gaussian fit to find the peak center (fig. S2b). We repeat this procedure for each acquired frame and take the mean value for x/y as the pixel center and line center in the image frame. fig. 2c-d show the distribution of Gaussian peaks.

##### 1.1.3 Pixel Size Determination

The image pixel size must be known to convert the pixel positions from pixel-space to stage-space. The pixel size was measured using a field of 5 μm microspheres (Bangs Laboratories, Inc, no. FCFR008) imaged and then translated in 1 μm steps along a single axis of the piezoelectric stage after at least 1 frame-time of acquisition. fig. S3a shows the field of beads in image frame-space of frame 3. The fitted centroids were labelled and color-coded in red. The stage is translated and imaged each step and fig. S5b shows the resulting field at frame 91. The beads centroids color-coded in green are visibly shifted from their original green location, apparent when the two frames are overlaid in fig. S5C. The distance each bead moves along x axis between frames is $\Delta_{pixel}$ and the distance each bead moves along x axis between frames is $\Delta_{line}$. This process was repeated across the 75 µm range of the stage and performed twice for each axis. The locations of the microspheres in the frame space were tracked using a circular Hough Transformation and distance minimization algorithm to identify each bead in every frame. The relative change in image pixel or line centroid location versus the relative change in X or Y stage position yields a linear fit corresponding to the image size of the pixels in px/µm (fig. S5d-g). The slope direction for each axis is a function of the difference in orientation between the laser scan and the stage and the sign must be known and accounted for to convert the image from frame-space to stage-space.

##### 1.2 Sample Preparation

For these single virus tracking experiments, we incorporated fluorescent protein into VSV-G VLPs by fusing either eGFP, or miRFP670, to HIV-1 Vpr which is packaged within the pseudovirus nucleocapsid. 3D-TrIm trajectories were acquired on cells labeled with SYTO61 (targeted to nucleic acids) or SiR650-actin (targeted to f-actin). The cell labels were chosen to maximize chromatic separation between the tracking and imaging, and the largest contributor to tracking crosstalk was cell autofluorescence by single-photon excitation, which was low enough not to perturb the active-feedback single-virus tracking. We cultured monolayers of either HeLa, BJ fibroblasts, or LDLR-deficient GM701 fibroblasts on glass coverslips. Alternatively, to achieve multi-layered cells, we cultured HT29-MTX cells on inverted matrix support filters (*61*). This selection of cell types offered diversity in morphology, extracellular environment, and cell surface receptor concentration to observe their influences on the early stages of viral contacts.

###### 1.2.1 Vpr-mGreenLantern Plasmid Construction

The expression vector Vpr-mGreenLantern, used to generate internally mGreenLatnern labelled VSV-G VLPs, was constructed as follows. The vector backbone was derived from H2B-mGreenLantern (Addgene, no. 164464) and amplified by PCR with the following primers: 5′-GATCCACCGGTCGCCAC-3′ and 5′-CATGGTGGCGGTACCGTC-3′. DNA encoding for Vpr was amplified by PCR with the following primers: 5′-TCGACGGTACCGCCACCATGGAACAAGCCCCAGAAGACCA-3′ and 5′-ATGGTGGCGACCGGTGGATCGGATCTACTGGCTCCATTTCTTCTTGC-3′, using eGFP-Vpr (construction mentioned in main text) as the template. PCR products were overlapped and assembled to create Vpr-mGreenLantern with In-Fusion Snap Assembly cloning kits (Takara Bio, no. 638947); successful insertion was verified by sequencing.

###### 1.2.2 Antibody Labelling

To produce a labeled secondary antibody for immunofluorescence assays, 50 µg of goat anti-mouse IgG H&L (Abcam, #ab6708) was dialyzed against 1 L of 1× PBS at 4 °C for 4 h (D-tube mini, MWCO 6-8 kDa, Millipore Sigma, #71504-M). The volume of the recovered antibody was measured and to this 1/10 the volume of 100 mM $NaHCO_3$ (pH = 8.3) was added. Next, to initiate labeling 5-fold molar excess of AF555-NHS ester (ThermoFisher, no. A20009), dissolved and subsequently diluted in anhydrous DMSO, was added so that the final volume of DMSO was < 2%. The reaction was left to continue at room temperature for 1 h. The solution was applied to a desalting column (Zeba™ Spin, 7K MWCO, ThermoFisher, #89882) pre-equilibrated with 10 mM Tris-HCL, pH = 7.5, and 0.01% $NaN_3$, to terminate the reaction and remove uncoupled dye.

##### 1.2.3 Immunofluorescence

The packaging efficiency of Vpr.StayGold inside the VLP was analyzed by two different immunofluorescence assays, one targeted against the inner capsid the other against the external envelope glycoprotein. VLPs were adhered to autoclaved glass coverslips (VWR, # 48380-046) coated with poly-L-lysine (#P6282, Millipore Sigma), overnight in PBS at 4°C.

VLPs were first fixed for 20 min using 7.4% formalin (Fisher, no. BP531-500) in PBS. Subsequently, washed (3×, 5min) and permeabilized for 10 min in T-PBS (0.1% Triton X-100 in PBS (pH=7.2)). Coverslips were blocked with buffer containing 10% normal goat serum (MP Biomedicals, #IC19135680), 0.2 M Glycine, and 0.1% Triton X-100 in PBS (pH=7.2) for 90 min at room temperature.

VLPs were stained for capsid protein using mouse anti-HIV-1 p24 gag monoclonal antibody (The following reagent was obtained through the NIH HIV Reagent Program, Division of AIDS, NIAID, NIH: Anti-Human Immunodeficiency Virus 1 (HIV-1) p24 Gag Monoclonal (#24-3), ARP-6458, contributed by Dr. Michael Malim), at 2 ng $\mu L^{-1}$ in blocking buffer 120 min at room temperature. Coverslips were washed (3×, 5 min) in T-PBS to remove any unbound primary antibody.

VLPs were stained for VSV-G envelope glycoprotein using mouse anti-VSV-G monoclonal (Kerfast, #EB0010), at 2 ng $\mu L^{-1}$ in blocking buffer 120 min at room temperature. Coverslips were washed (3×, 5 min) in T-PBS to remove any unbound primary antibody.

In both scenarios as a secondary antibody, AF555-labelled Goat Anti-Mouse IgG (described above) was used at 4 ng $\mu L^{-1}$ in blocking buffer for 120 min at room temperature. Finally, coverslips were washed with T-PBS (3×, 5 min), and mounted in PBS. In control experiments the same procedure was followed except either the primary or secondary antibody incubation was omitted.

Immunostained VLPs were imaged on a spinning disk confocal (Andor Dragonfly 505) on a Leica DMi8 inverted microscope using 100x/1.40-0.70 HCX PL APO (Leica 11506210) oil objective and 488 nm and 561 nm laser lines for excitation (40 µm pinhole). Images were captured on an Andor iXon Life 888 1024×1024 EMCCD camera, and the system was controlled by Fusion 2.0. (Duke University Light Microscopy Core Facility NIH Shared Instrumentation grant 1S10RR027867-01).

Intensity based colocalization was performed using IMARIS software (Oxford Instruments) to extract the Pearson's coefficient for each field of view. Independently, Vpr.StayGold centers were identified in MATLAB (MathWorks) by determining the intensity maximum of each foci and the intensity at the corresponding position in the AF555 channel.

##### 1.2.4 Infection Assay

To assess the infectivity of VSV-G pseudotyped VLPs containing fluorescent Vpr, 297T/17 cells were inoculated with VSV-G lentiviral vector encoding the ZsGreen1 reporter construct pLVXS-ZsGreen1-Puro. 293T/17 cells (Duke Cell Culture Facility, ATCC# CRL-11268) were grown using complete DMEM which comprised of DMEM Media (Corning, no. 10-013-CV) supplemented with 10% FBS (Millipore Sigma, no. F2442), and 1 × penicillin-streptomycin (Corning, #30-002-CI) in T12.5 flask (Genesee Scientific, no. 25-205). Cells were plated in complete DMEM at 5×104 cells/well in an 8 well μ-slide, glass bottom (Ibidi, no. 80827). 297T/17 cells were immediately inoculated with a multiplicity of infection (MOI) of 1, 10, or 100 transfecting units per cell of lentiviral vector. The infected cultures were incubated at 37 °C in an atmosphere of 5% CO2.

At 72 h p.i., cells were fixed with 4% paraformaldehyde (PFA) solution in 1 × PBS (Ph = 7.2) for at least 20 minutes. Then the cells were stained with 1.5 μM of Hoechst 34580 (ThermoFisher, no. H21486) diluted with 1 × PBS. The fixed sample was kept at 4 °C until imaging within 3 days.

The imaging was conducted on a spinning disk confocal microscope (Andor Dragonfly 505), equipped with 40x/1.3 HC PL APO CS2 (Leica 11506358) oil objective, and 405 nm and 488 nm laser lines for excitation (40 μm pinhole). Simultaneous two-color imaging was performed at 2.5 % and 0.5 % 405 nm and 488 nm laser power, respectively. Images were captured on an Andor iXon Life 888 EMCCD camera with 200 msec exposure time for both colors.

Nuclei and cell boundaries of ZsGreen1-positive cells were identified using the CellProfiler image analysis software. The thresholds were obtained from the negative control group with 95% confidence.

##### 1.2.5 Coverslip Poly-L-lysine Coating

The poly-L-lysine solution was prepared by adding 50mL of sterile tissue culture grade water to a 5 mg of poly-L-lysine (Millipore Sigma, no. P6282, desiccated solid stored at -20 °C). Once the solution was prepared, it could be stored at 4 °C and reused up to three times.

Autoclaved glass coverslips (VWR, no. CLS-1760-025) were autoclaved and transferred into a 6-well plate. The coverslips were first coated with 1.5mL EtOH (1% HCl v/v) and left at room temperature for 5 minutes. The solution was removed, and the coverslips were dried under 60 ℃ for 20 minutes. After cooling down to room temperature, the coverslips were coated with 1.5mL 0.5mg/mL poly-L-lysine. To ensure even coating of the culture surface, the 6-well plate was rocked gently. After 5 minutes, the solution was removed, and coverslips were thoroughly rinsed with sterile tissue culture grade water for at least twice. The coverslips were dried in clean bench hood for at least 2 hours before introducing media and samples. Plates were stored at 4 ℃ until use.

##### 1.2.6 Autofluorescence Control Experiment

To exclude the influence of autofluorescence from 293T/17 cells, a control experiment was executed by performing SVT on 293T/17 with the absence of any VLPs. The other experimental conditions remained the same as mentioned in main text - Materials and Methods – Microscope

Sample Preparation. The tracking initialization threshold is 22 kHz for experiments with VLPs. While for control experiments without fluorescent particles, the tracking initialization threshold is tuned down to 16 kHz to be able to collect an adequate number of trajectories.

The results of control experiments are shown in fig. S11. Autofluorescence does affect tracking to some degree but could be identified and filtered based on the pick-up intensity and tracking duration. Any trajectory with pick-up intensity lower than 21 kHz (mean + 2SD, 95% confidence interval) was considered as autofluorescent. The durations of trajectories tracking autofluorescent puncta was fitted into Burr distribution model and the 95% confident bound was identified as 240 second.

1.3 Data Analysis

The following sections describe protocols used to quantitatively analyze simultaneously acquired VLP trajectories and live cell imaging data.

1.3.1 Analysis of VLP Size by 3D SMART

The size and brightness of Vpr.StayGold or Vpr.eGFP incorporated VLPs was evaluated by real-time 3D tracking to extract the diffusion coefficient and particle emission rate. Free virus particles were tracked in HEPES pH = 7.4 buffered solution (live cell imaging solution, ThermoFisher, no. A14291DJ) at room temperature. The tracking microscope configuration was identical to that used for tracking and imaging as described in the Methods section. The average excitation power of the 488 nm laser was ~40 nW at the focus.

1.3.2 Mean Square Displacement Analysis

The diffusion coefficient can be obtained by linear fitting of the mean square displacement (MSD) with lag time (τ). The MSD was calculated using the definition:

$$MSD(\tau) = N^{-1}\Sigma_{n=1}^{N}\left((x(n+\tau) - x(n))^2) + (y(n+\tau) - y(n))^2) + (z(n+\tau) - z(n))^2)\right) \quad (3)$$

Here, $x(n)$, $y(n)$, and $z(n)$ are the coordinates of the trajectory at timepoint $n$. $N$ is the total number of data points.

The power exponent, alpha (α), was calculated from the power law fit of MSD(τ).

$$MSD(\tau) = 2nD\tau^{\alpha} \quad (4)$$

Here, $n$ is the number of dimensions, $D$ is the diffusion coefficient and $\alpha$ is the power exponent used to determine superdiffusion and subdiffusion.

For single virus tracking only experiments, without cells, the hydrodynamic radius ($r$) of the particles was calculated using the Stokes-Einstein relation:

$$D = \frac{k_B T}{6\pi\eta r} \quad (5)$$

Here, $k_B$ is the Boltzmann constant, $T$ is temperature, and $\eta$ is viscosity of solution.

##### 1.3.3 Determination of Emission Intensity of a Single Labeled Vpr

The emission intensity of a single Vpr labelled with eGFP or StayGold was determined by photobleaching step analysis. Photobleaching experiments were performed on fixed VLPs packaging Vpr-eGFP and Vpr-StayGold. VLPs were adhered to autoclaved quartz coverslips (EMS, no. 72256-06) coated with poly-L-lysine (Millipore Sigma, no. P6282) coated coverslip overnight in PBS at 4 °C. Fixed VLPs packaging fluorescent Vpr were held in the laser focus by engaging the tracking system. For photobleaching experiments, the 488nm laser power was ~ 200 nW at the objective focus. The intensity level of a single eGFP or StayGold chromophore was established by tracking the VLP until the intensity reached the background level.

The final step of photobleaching was identified and validated by performing a sliding window *t*-test. The window size is 5 seconds, and the slide step size is 0.1 seconds. The p-values were plotted over time and the timing of smallest p-value was used to determine the critical final photobleaching step.

#### 2 Figures and Movies

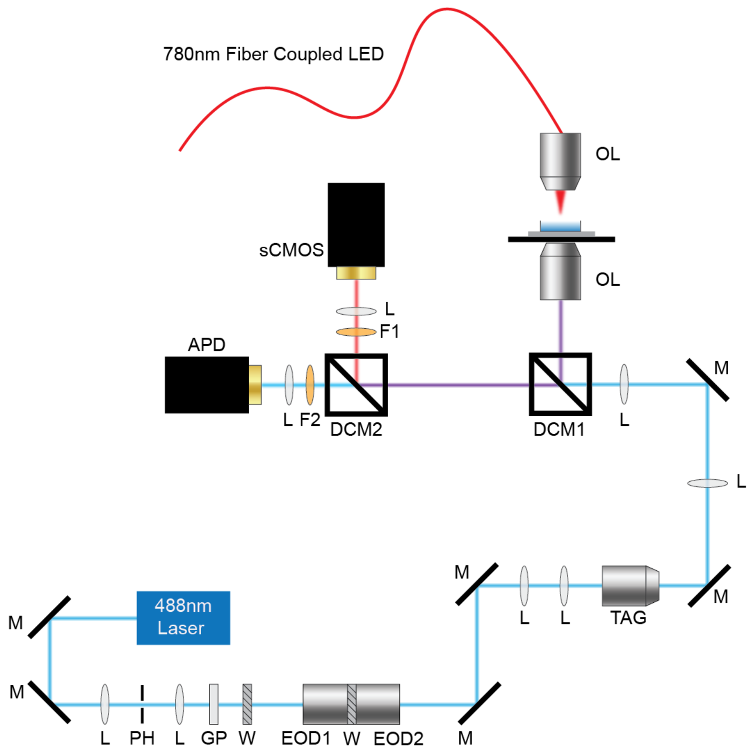

**Fig. S1. Instrument diagram**

M: mirror, L: lens, PH: pinhole, GP: Glan-Thompson Polarizer, W: half wave plate, EOD: electro-optic deflector, TAG: tunable acoustic gradient lens, DCM: dichromatic mirror, OL: objective lens, F: fluorescence emission filter.

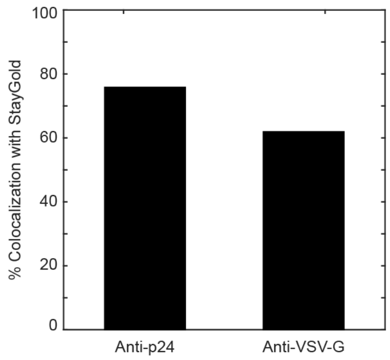

**Fig. S2. Colocalization efficiency with StayGold, related to Fig. 1b-c**

The colocalization efficiency of anti-p24 and anti-VSV-G with StayGold signal is 75.8% and 61.9%, respectively. The red signal in Fig. 1b-c corresponds to capsid (anti-p24) or envelop protein (anti-VSV-G), while the green signal corresponds to StayGold. The colocalization efficiency was calculated by dividing the number of particles with colocalized red and green signals by the sum of colocalized and green-signal only particles (corresponding to the total amount of 'trackable' particles).

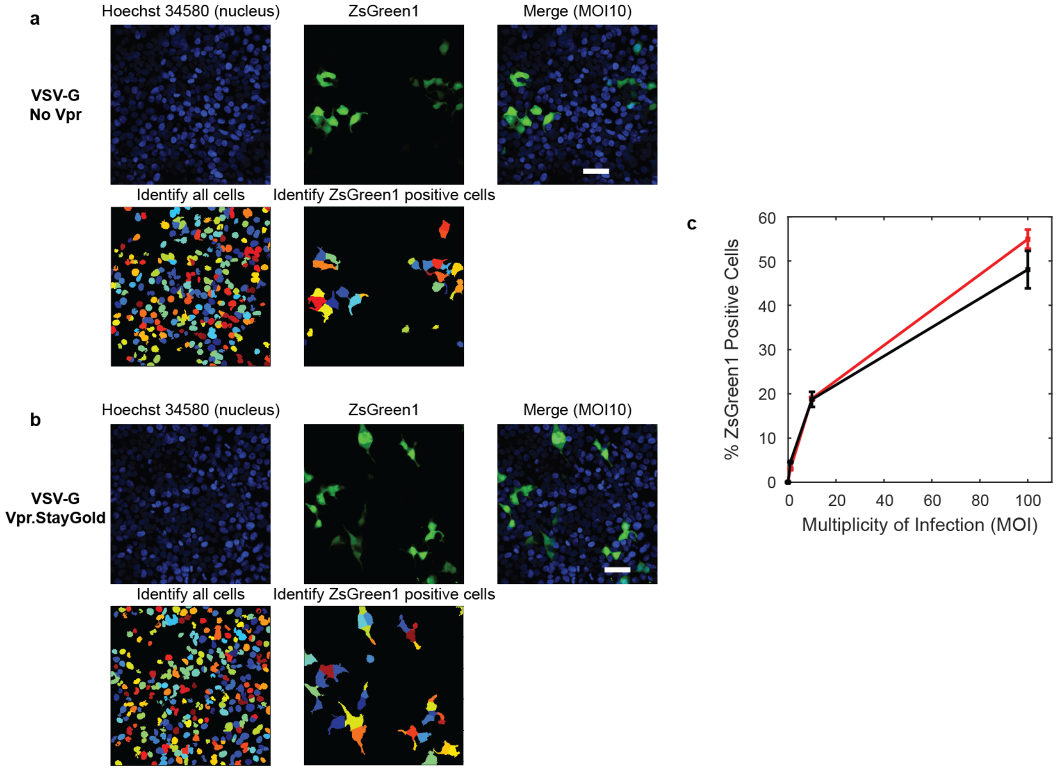

**Fig. S3. VSV-G gene delivery remains with the presence of Vpr-StayGold**

(**a**) ZsGreen1 expression in 293T/17 cells after infection with VSV-G carrying ZsGreen1-reporter gene without encapsulated Vpr. (**b**) ZsGreen1 expression in 293T/17 cells after infection with VSV-G carrying ZsGreen1-reporter gene with encapsulated Vpr-StayGold. 293T/17 cells were stained with Hoechst 34580, 72h post-transduction at MOI 10 (top panel). The total number of cells and number of cells expressing ZsGreen1 were detected using CellProfiler analysis software (bottom panel) over 3 independent experiments. Scale bar = 50 µm. (**c**) Average ± SEM ($n \approx 3 \times 10^4$ cells) of the percentage of cells expressing ZsGreen1 after transduction with VSV-G No Vpr (red) or VSV-G Vpr-StayGold (black).

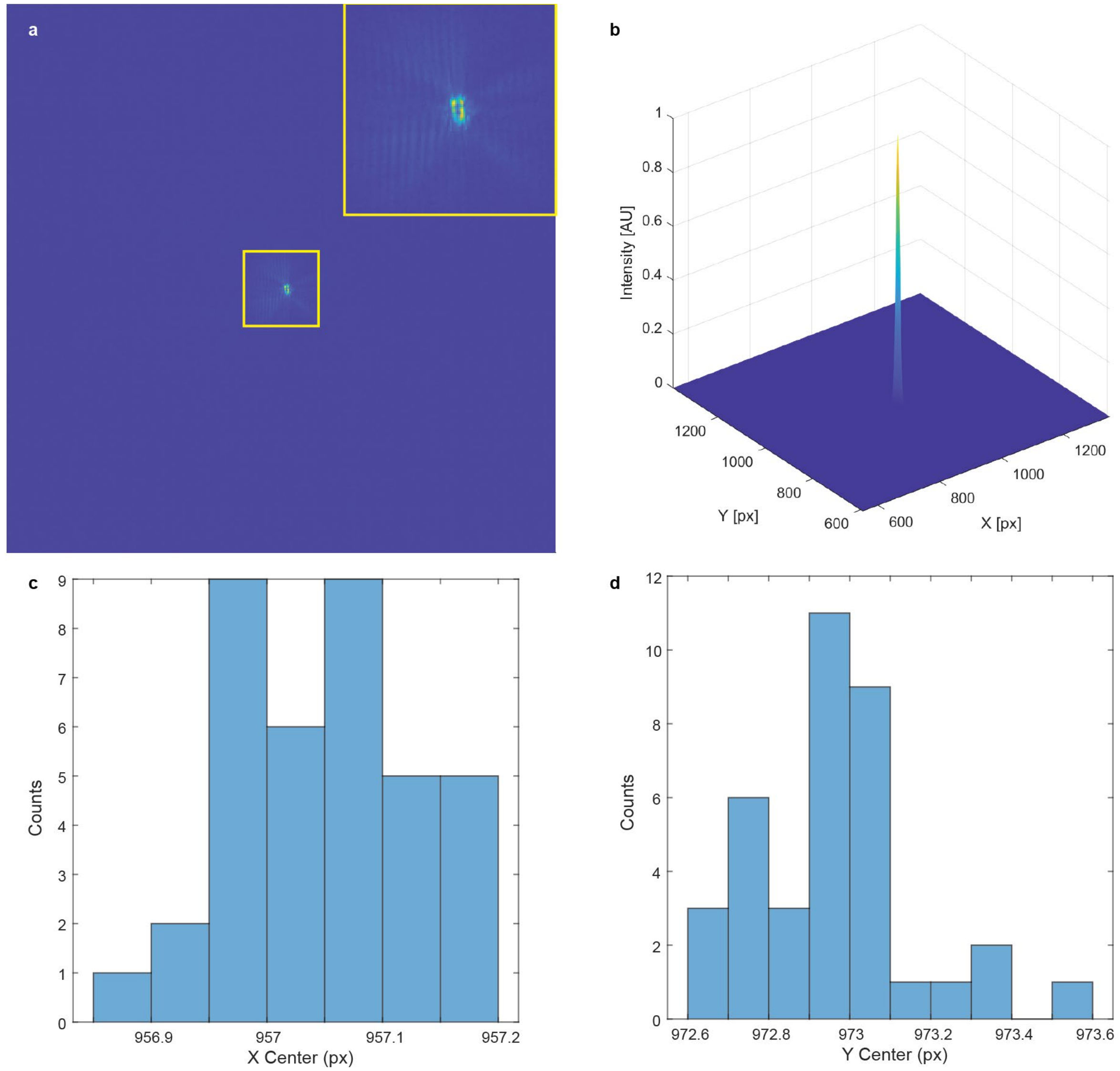

**Fig. S4. Track center calibration**

Track center registration calibration. (**a**) sCMOS image of the 488-nm tracking laser. Insert shows magnified view. (**b**) 2D Gaussian fitting of intensity is used to find track center location. (**c**) Histogram of X centroid coordinates. (**d**) Histogram of Y centroid coordinates. Histograms in (**c**) and (**d**) are from n = 3 independent experiments.

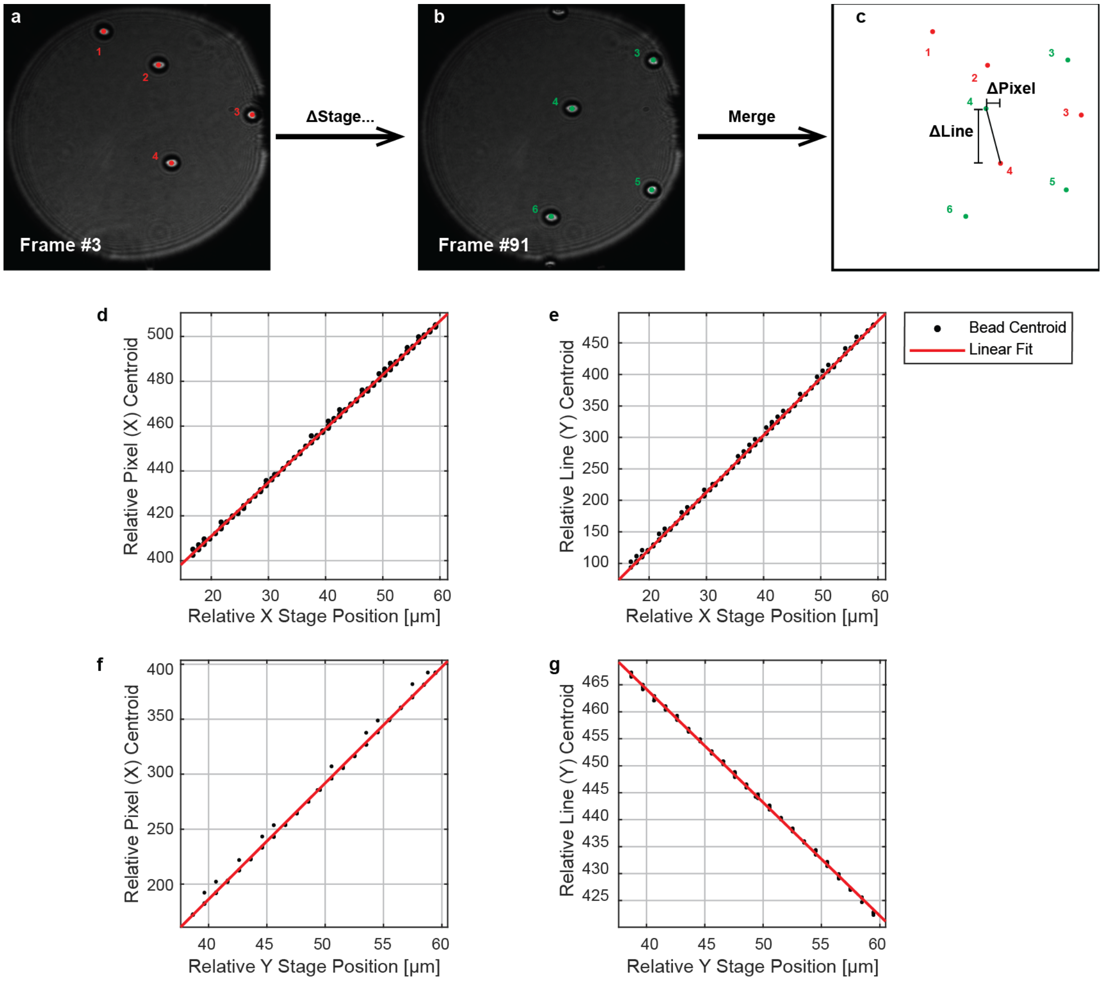

**Fig. S5. Pixel size calibration**

Track center registration calibration. (**a**) Frame #3 of a field of 5 μm microspheres. The centroids of microsphere are labelled and color-coded in red. (**b**) After a few steps of piezoelectric stage movement in single direction, frame #91 shows significant shifting of beads' position. The centroids are labelled and color-coded in green. (**c**) Overlay of frame #3 and #91. (**d** and **e**) Plot of relative pixel and line displacement versus stage displacement along X axis. (**f** and **g**) Plot of relative pixel and line displacement versus stage displacement along Y axis.

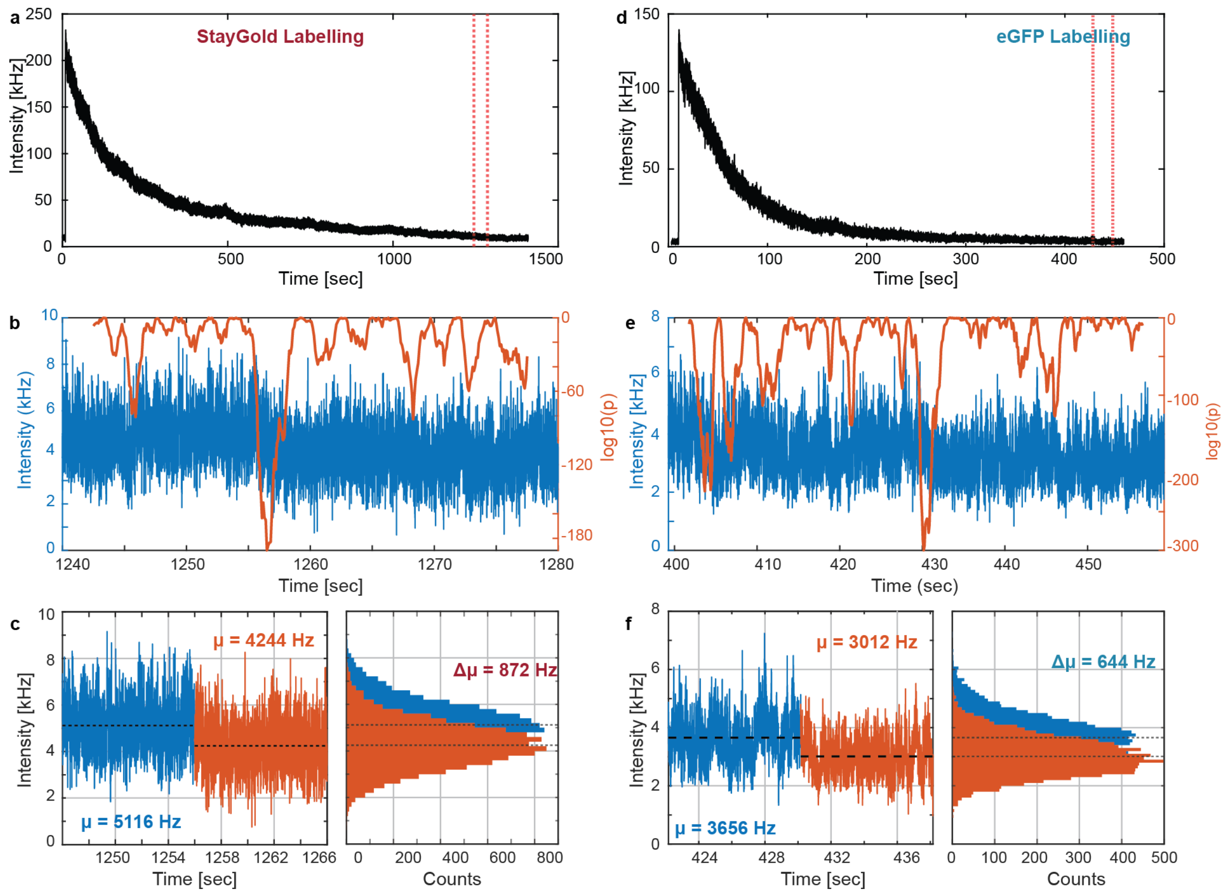

**Fig. S6. Determination of the emission intensity of a single Vpr-StayGold and Vpr-eGFP**

(**a**) Intensity trace of a Vpr-StayGold VLP immobilized on a glass coverslip. (**b and e**) Sliding window *t*-test was performed on a segment of the intensity trace to determine the last step of photobleaching for Vpr-StayGold and eGFP-StayGold respectively. The timing of the smallest p value was identified as the change step. (**c, left**) Zoom in on the final bleaching step, with the intensity before the bleach shown in blue and the background intensity shown in orange. (**right**) Histogram of the intensities from **left**, showing the magnitude of the final bleaching step to be 872 Hz for a single Vpr-StayGold. (**d**) Intensity trace of an Vpr-eGFP VLP immobilized on a glass coverslip. (**f, left**) Zoom in on the final bleaching step, with the intensity before the bleach shown in blue and the background intensity shown in orange. (**right**) Histogram of the intensities from **left**, showing the magnitude of the final bleaching step to be 644 Hz for a single Vpr-eGFP.

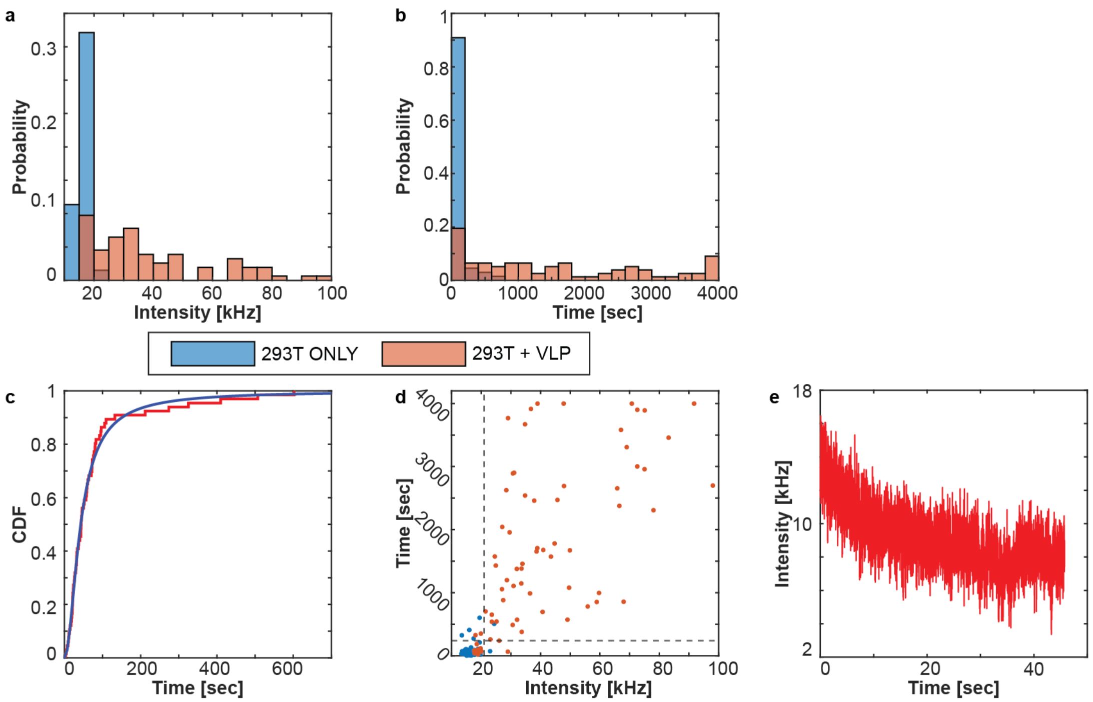

**Fig. S7. Control experiment of tracking in 293T/17 cells without VLPs**

(**a**) Histogram of pickup intensity of SVT with and without the presence of StayGold-VLPs (n = 66 and 77 respectively). (**b**) Histogram of tracking duration of SVT with and without the presence of StayGold-VLPs. (**c**) The cumulative distribution function (CDF) of tracking duration without VLPs (red) overlayed with fitting result to Burr distribution (blue). (**d**) Scatter plot of pickup intensity and tracking duration of SVT with and without the presence of StayGold-VLPs. The dashed lines showed the 95% confidence bound for the pickup intensity and tracking duration of autofluorescent puncta respectively. (**e**) An example intensity trace of autofluorescent puncta showed low pickup intensity and prompt photobleaching.

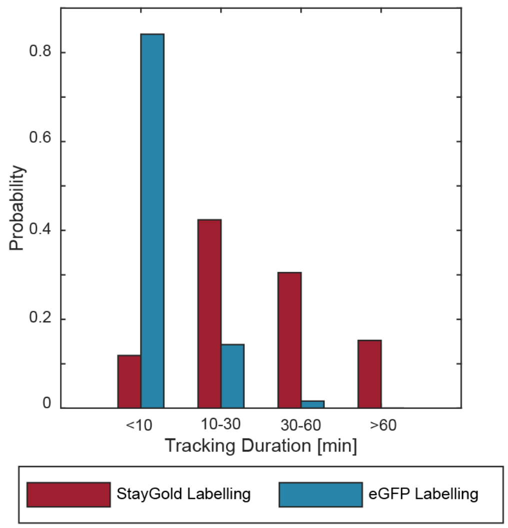

**Fig. S8. StayGold labeling significantly extends observation time**

The observation period of most trajectories of eGFP-VLPs (blue) remains within 10 minutes. With StayGold labelling and the enhanced photostability, the tracking duration was extended and a considerable amount of StayGold-VLPs (red) can be tracked for over an hour.

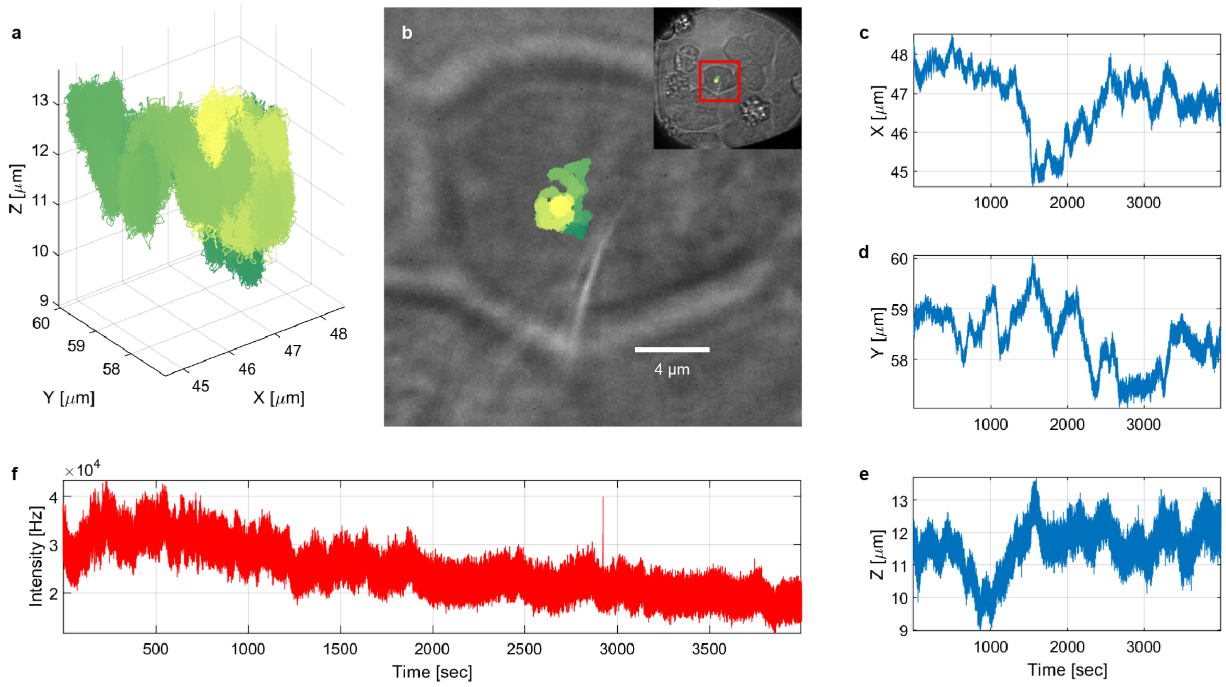

**Fig. S9. An example of tracking VLP-StayGold for over an hour**

(**a**) A 3D trajectory over an hour long of tracking a VSV-G lentiviral particle packaging Vpr-StayGold in a live 293T/17 cell. (**b**) bright field image overlapped of the top-down view trajectory in (**a**) obtained during the collection of the trajectory by 3D-SMART. Insert: larger view of cell. (**c-f**) X, Y, Z, and intensity trace of the trajectory in (**a**).

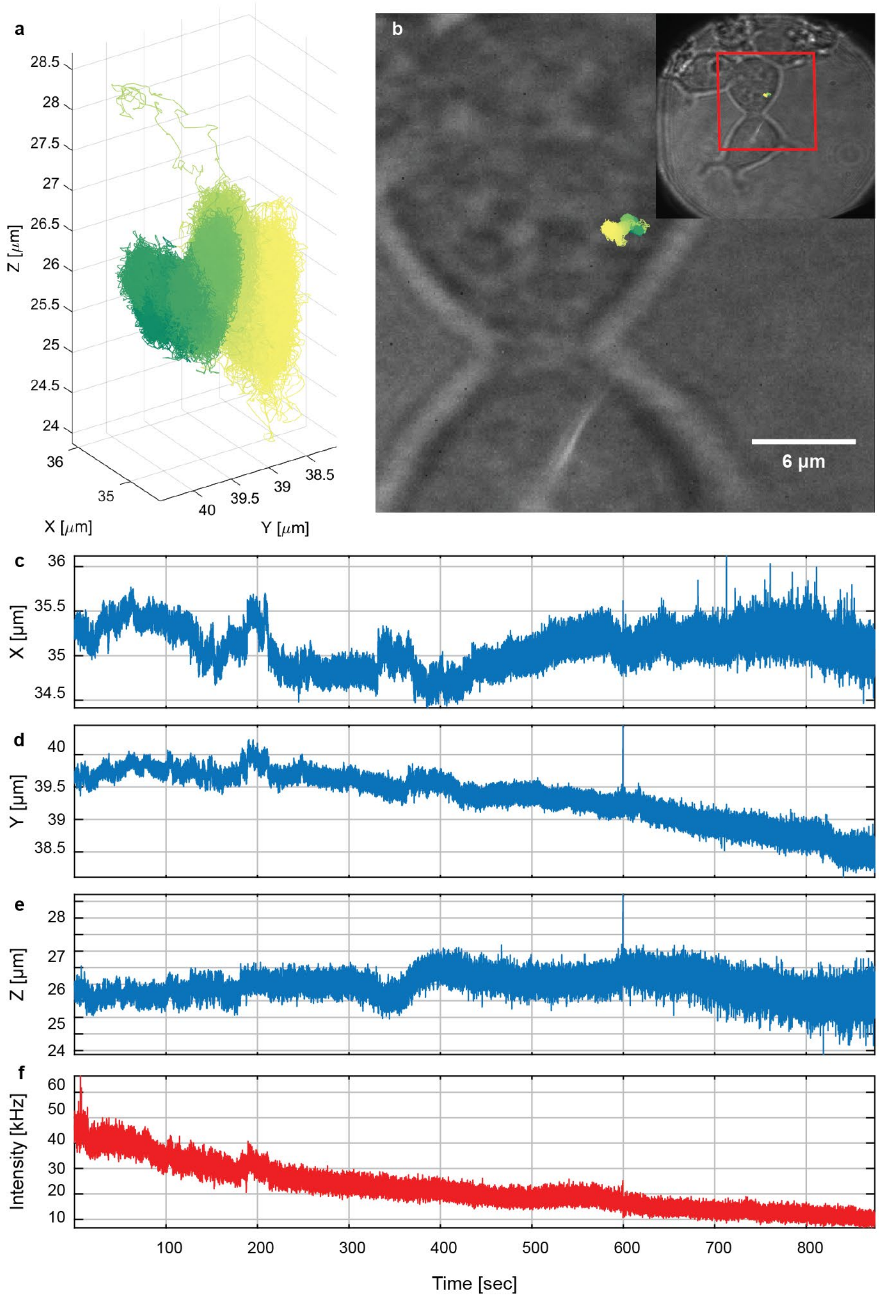

**Fig. S10. Example trajectory of VSV-G Vpr-eGFP, related to Fig. 5 and fig. S6**

(**a**) 3D trajectory of a VSV-G lentiviral particle packaging Vpr-eGFP in a live 293T/17 cell. (**b**) bright field image overlapped of the top-down view trajectory in (**a**) obtained during the collection of the trajectory by 3D-SMART. Insert: larger view of cell. (**c**-**f**) X, Y, Z, and intensity trace of the trajectory in (**a**).

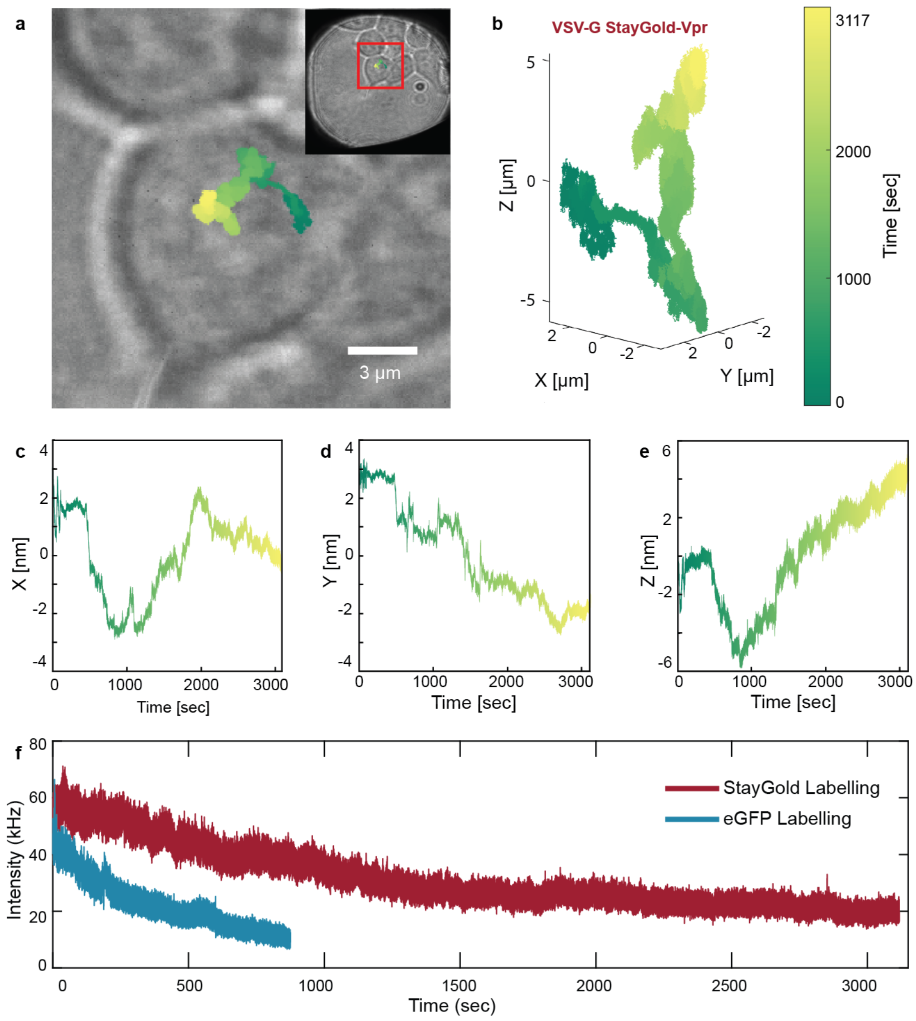

**Fig. S11. Complete trajectory of VSV-G Vpr-StayGold VLP fit to spherical surface, related to Fig. 6**

(**a**) bright field image overlapped of the top-down view trajectory in (**b**) obtained during the collection of the trajectory by 3D-SMART. Insert: larger view of cell. (**b**) 3D trajectory of a VSV-G lentiviral particle packaging Vpr-StayGold in a live 293T/17 cell. (**c-e**) X, Y and Z trace of the trajectory in (**b**). The trajectory and XYZ traces of (**a-e**) are color-coded by time with the same colormap in (**b**). (**f**) Raw intensity trace of the trajectory in (**b**) and a trajectory of a VSV-G lentiviral particle expressing Vpr-eGFP in a 293T/17 cell (see fig. S7) shows better photostability and longer observation time of StayGold labelling.

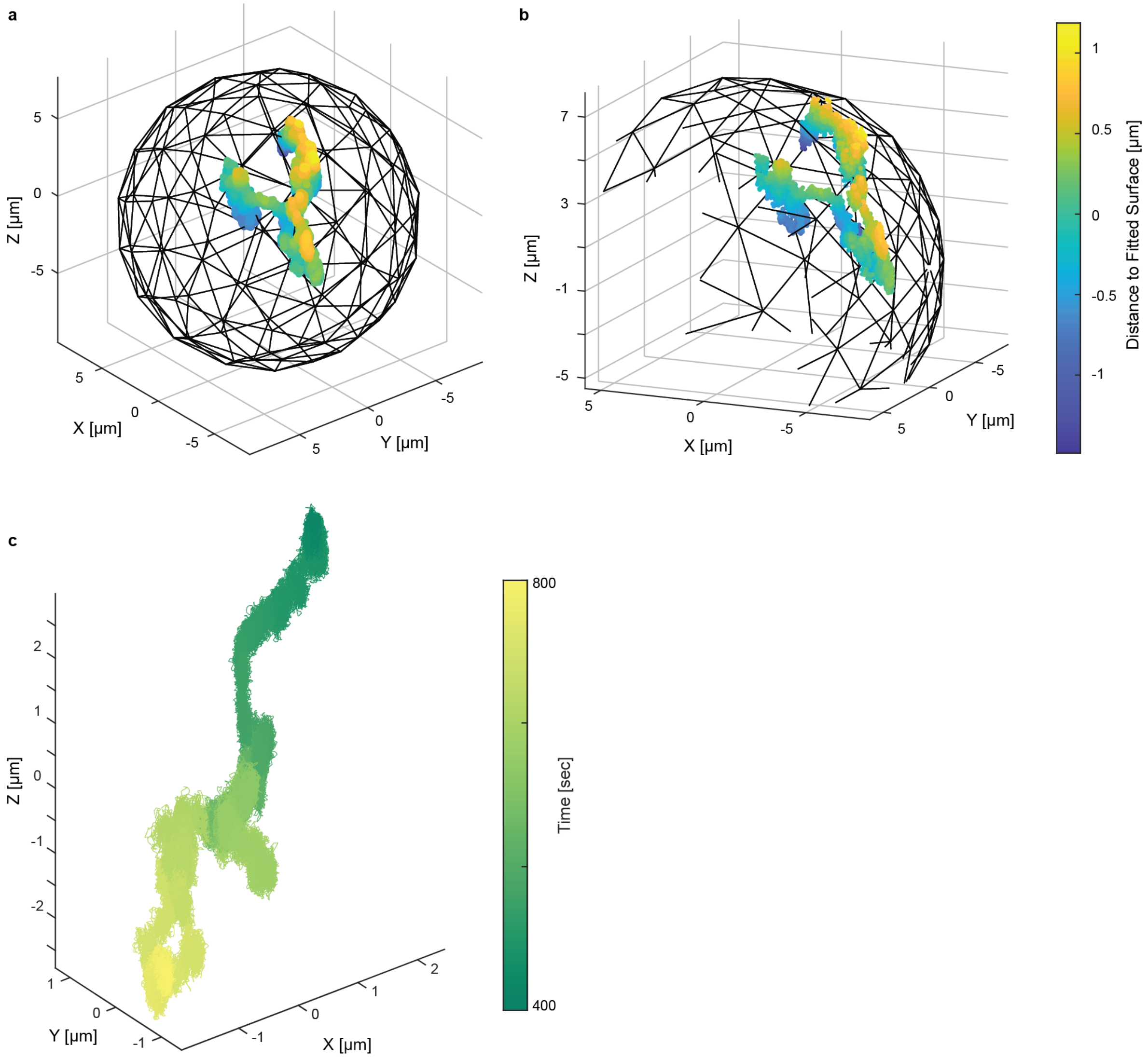

**Fig. S12. Viral trafficking on a spherical surface and comparison to typical eGFP-VLPs lifetime, Related to Fig. 6**

(**a**) The segment of trajectory (fig. S6, first ~2000 sec) overlayed with the fitted sphere with a diameter of ~ 8.6µm in the cartesian coordinate. The segment was color-coded by the distance of each point to the fitted surface to visualize the goodness of the fitting. (**b**) A different view of (**a**) from another angle to show the curvature of the segment. (**c**) A 400 second segments from trajectory showed in Fig. 6 to mimic typical tracking duration of eGFP labelled VLPs.

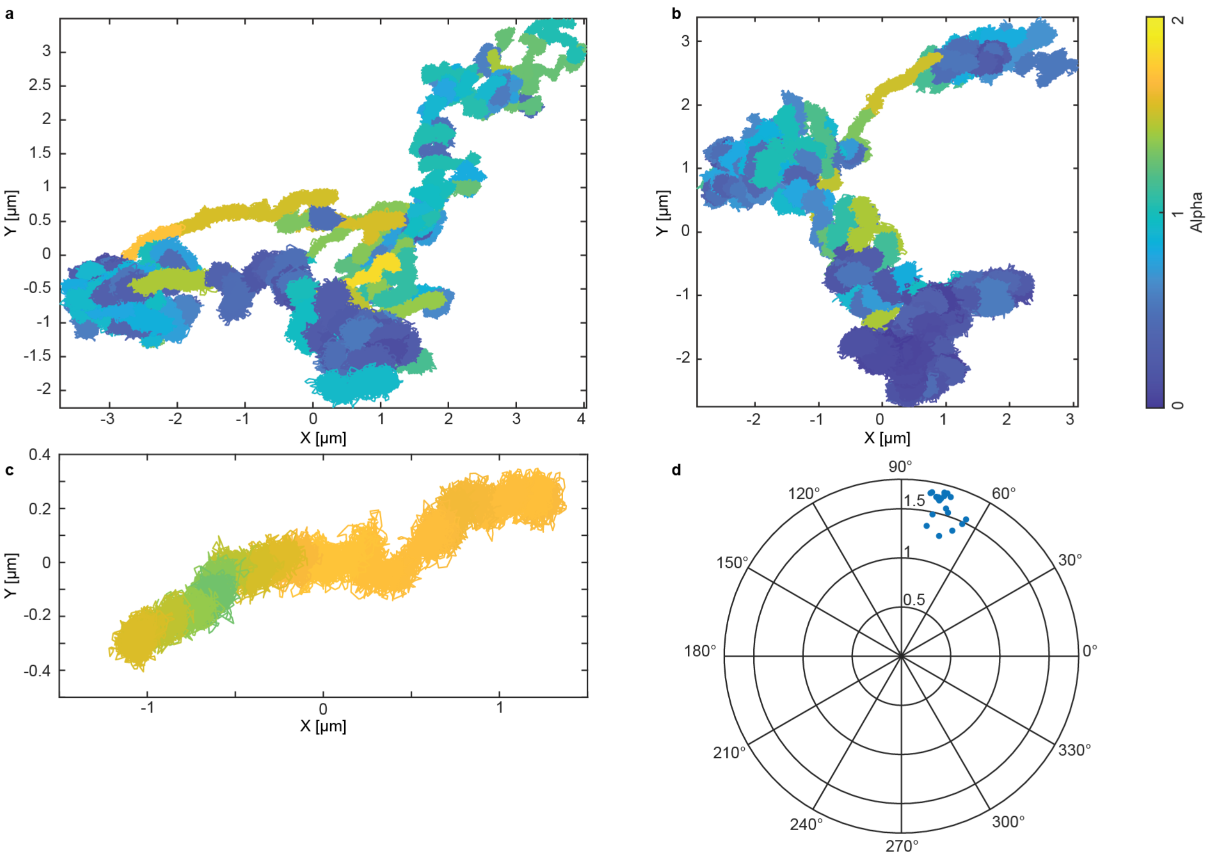

**Fig. S13. MSD analysis shows difference between active trafficking and membrane diffusion** (**a-b**) *XY* projection of trajectories in Fig. 5 (**a**, ~50 minutes, linear trafficking) and Fig. 6 (**b**, ~35 minutes, membrane diffusion, see also fig. S6) color-coded by alpha. (**c**) 40-seond segment from (**a**) with a low calculated diffusion coefficient of 0.0668 ± 0.0068 $\mu m^2/sec$ but high alpha 1.58 ± 0.13. (**a-c**) shared the same color bar. (**d**) Polar scatter plot of the drift velocity direction and alpha calculated from the segment in (**c**).

**Movie S1. Tracking VSV-G Vpr-StayGold VLP in live 293T/17 cells, related to Fig. 5.**

Movie S1. Tracking VSV-G Vpr-StayGold VLP in live 293T/17 cells, related to Fig. 5.
(Left) Trajectory (~ 50 minutes) was color mapped by time with a sampling rate of 1 kHz. (Right top) Brightfield imaging of unlabeled live 293T/17 cells overlayed with XY projected virus trajectory. Cells were color-coded by image intensity. (Right bottom) Intensity traces of two trajectories of VLPs labeled with StayGold (red, corresponding to the left trajectory) and eGFP (blue, fig. S7) respectively. The playback rate was constant 60 ×. Trajectories shared the same colormap and sampling rate. The intensity traces were synchronized with the trajectories and had the same playback rate.

**Movie S2. 3D reconstruction of viral diffusion on a curved cellular surface, related to Fig. 6.**

(Left) The segment of trajectory (~ 35 minutes) was color mapped by time with a sampling rate of 1 kHz. At ~ 8.5 second of the movie, the fitted spherical surface showed up and overlayed with the 3D trajectory. (Right) Brightfield imaging of unlabeled live 293T/17 cells overlayed with XY projected virus trajectory. Cells were color-coded by image intensity. The playback rate was constant 60 ×. Trajectories shared the same colormap, sampling rate and playback rate.
